# Conserved protein sequence-structure signatures identify antibiotic resistance genes from the human microbiome

**DOI:** 10.1101/2025.10.01.679039

**Authors:** L. Bartrop, E. Beauchemin-Lauzon, F. Grenier, S. Rodrigue, L.P. Haraoui

## Abstract

Bacteria exhibiting antimicrobial resistance (AMR) is a problem that has grown to become a significant public health challenge worldwide. Antibiotic resistance genes (ARGs), determinants of AMR, mostly emerge from non-clinical settings. Identifying previously undetected ARGs in the human microbiome which confer resistance to clinical concentrations of antibiotics is a crucial component of addressing AMR, yet can be hindered by their low homology to existing ARGs. Here, we attempt to address this by focussing on functionally important protein regions. ARG-PASS (ARG-PAirwise Sequence vs Structure) represents a novel protein function prediction method which leverages a one-class support vector machine trained on pairwise primary and tertiary distributions of structurally conserved regions of proteins encoded by ARGs. ARG-PASS was applied to six reference strains of the Human Microbiome Project. Nine candidates were selected for experimental verification and all were functionally confirmed when expressed in *E.coli*, belonging to ARG classes: APH, *dfr*, class B and C β-lactamases, and penicillin binding proteins. We also used ARG-PASS directly on protein structures within the AlphaFold database and predicted a *phnP* gene (metallo-β-lactamase fold), which is highly divergent from existing β-lactamases and had activity against ampicillin. In total, 80% of the tested genes confer resistance at CLSI resistant breakpoints and the remainder represent ‘pre-resistance’ genes, with activity but not at clinically relevant minimum inhibitory concentrations. We suggest pre-resistance genes may preferentially evolve into clinically relevant resistance determinants. ARG-PASS represents a novel and precise method of identifying previously uncharacterized ARGs from DNA databases, contributing to resistance surveillance and antibiotic stewardship.

## Background

The enthusiasm following the discovery of antibiotics was rapidly overshadowed by the rise of bacteria exhibiting antimicrobial resistance (AMR), defined as the ability of microbes to survive treatment with antibiotics that once effectively controlled them. AMR has since become one of the most pressing global public health challenges of our time (O’Neill, 2016). Antibiotic resistance genes (ARGs), key drivers of AMR, predate the use of antibiotics (D’Costa et al., 2011). For the most part, they are present in environmental bacteria but can be selected in response to anthropogenic antibiotic exposure, and subsequently pose a risk to human health once stably integrated into the human microbiome. This can occur through one of three main routes: (i) if they evolved in generalist or opportunist species which can also inhabit human microbiome niches; (ii) via horizontal gene transfer from environmental bacteria to human-associated bacteria by mobile genetic elements (MGEs); or (iii) by *de novo* evolution within the human microbiome (Crits-Christoph et al., 2022; Larsson & Flach, 2022). Previously unidentified (novel) ARGs can also evade detection by computational algorithms designed for their identification, due to their low homology to existing ARGs. As such, identifying novel ARGs from human microbiomes that confer clinical levels of resistance to antibiotics is a crucial component of addressing AMR for accurate surveillance, diagnostics, and antibiotic stewardship.

Existing computational tools for ARG discovery can be broadly categorised along two primary axes: (i) dependence on amino acid (aa) sequence similarity and (ii) strength of experimental (and clinical) validation. Traditional methods for identifying novel ARGs rely on a nearest neighbour approach using aa sequence alignment, typically with high seqID cutoffs (>60%) against databases of known ARGs (Alcock et al., 2023; Zhang et al., 2021). Profile hidden Markov models (HMMs) extend this capability through probabilistic modelling of protein classes (Feldgarden et al., 2019; Gibson et al., 2015; Yin et al., 2018). Experimental (wet-lab) validation of predictions (Berglund et al., 2019), facilitates their deposition into well-curated ARG databases, such as the Comprehensive Antibiotic Resistance Database (CARD) (Alcock et al., 2023). However, the amount of microbial proteins with homology to ARGs increases with decreasing seqID and ARGs can often share lower seqID. For example, at the time of discovery *bla*_NDM-1_ and *sul4* shared the highest seqID of ∼33% to *bla*_VIM_ and *sul2,* respectively (Yong et al., 2009; Razavi et al., 2017). Although effective for detecting close homologs, these approaches aren’t as suited for the discovery of more divergent sequences supported by experimental and clinical validation (Marathe et al., 2019).

When trained on the expanding volume of biological sequence data, ML-based classifiers have emerged as a potential alternative and/or complementary computational approach. Strong predictive performance has been reported with precisions exceeding 85% (Abbas et al., 2025; Arango-Argoty et al., 2018a; Hamid & Friedberg, 2020; Naik et al., 2025; J. Wu et al., 2023). However, ML-based classifiers will also favour the detection of less divergent ARGs if they are trained and/or validated on databases which ultimately suffer the same inherent biases, such as ARGs with little sequence variation, only from clinical settings, or without experimental support (Arango-Argoty et al., 2018a; Hamid & Friedberg, 2020; J. Wu et al., 2023). Whilst performance is high on internal validation, their generalisability to new sequences from other sources remains less clear.

Efforts to address this limitation expand training beyond sequences (Ghosh et al., 2025; Rannon et al., 2025). Pairwise protein sequence vs (experimental) protein structure distributions revealed structures are more conserved than their sequences (Illergård et al., 2009), potentially representing a different platform for ARG discovery. Pairwise comparative modelling (PCM) integrates protein structures encoded by ARGs and predicted novel ARGs from the intestinal resistome with an average seqID of 29.8%, and was supported by experimental validation (Ruppé et al., 2019). Nonetheless, without comparison of minimum inhibitory concentrations (MICs) to clinical breakpoints, clinical relevance of predictions is difficult to infer.

Since then, rapid developments in computational protein structure prediction tools (Jumper et al., 2021; R. Wu et al., 2022), has enabled progress in terms of accuracy (now in line with experimental solutions), and the number of structures available (>200 million predicted by AlphaFold2) (Varadi et al., 2024). Protein structural similarity can be quantified using the template modelling (TM)-score, which ranges from 0 (no structural similarity) to 1 (a perfect structural match) (Zhang & Skolnick, 2004). Pairwise sequence vs (computational) protein structure distributions across the microbial protein landscape provide further support for protein structure conservation relative to protein sequence (Koehler Leman et al., 2023).

A nearest-neighbour pairwise analysis (using the highest shared seqID per antibiotic resistant protein (ARP) structure and its corresponding TM-score), reveals ARP structures across well-known ARG classes exhibit coupling between sequence and structure, with an average median seqID of 82.9% and TM-score of 0.95 (Figure 1a and Figure 1b). This indicates that ARGs maintain both high sequence and structural conservation when compared to their closest homologue. However, ARGs can have high TM-scores despite lower seqID, with all classes containing representatives with a highest seqID below 50% (Figure 1a). For example, although sharing the highest seqID of ∼33%, ARP structures encoded by the aforementioned *sul4* and *bla*_NDM-1_ share TM-scores of over 0.9 to their respective closest homologues, encoded by *sul2* and *bla*_VIM_. A global pairwise analyses of the same dataset reveal lower seqIDs, with an average median of 30.5% and TM-score of 0.78, elucidating broader sequence and structure divergence among all representatives within each class (Figure 1c and Figure 1d). Importantly, existing computational tools potentially miss novel ARGs which have lower seqID to known ARGs. A computational approach which integrates the entire evolutionary context of the ARG class by a pairwise analysis, as opposed to the nearest-neighbour, may help to identify them.

**Figure 1.**
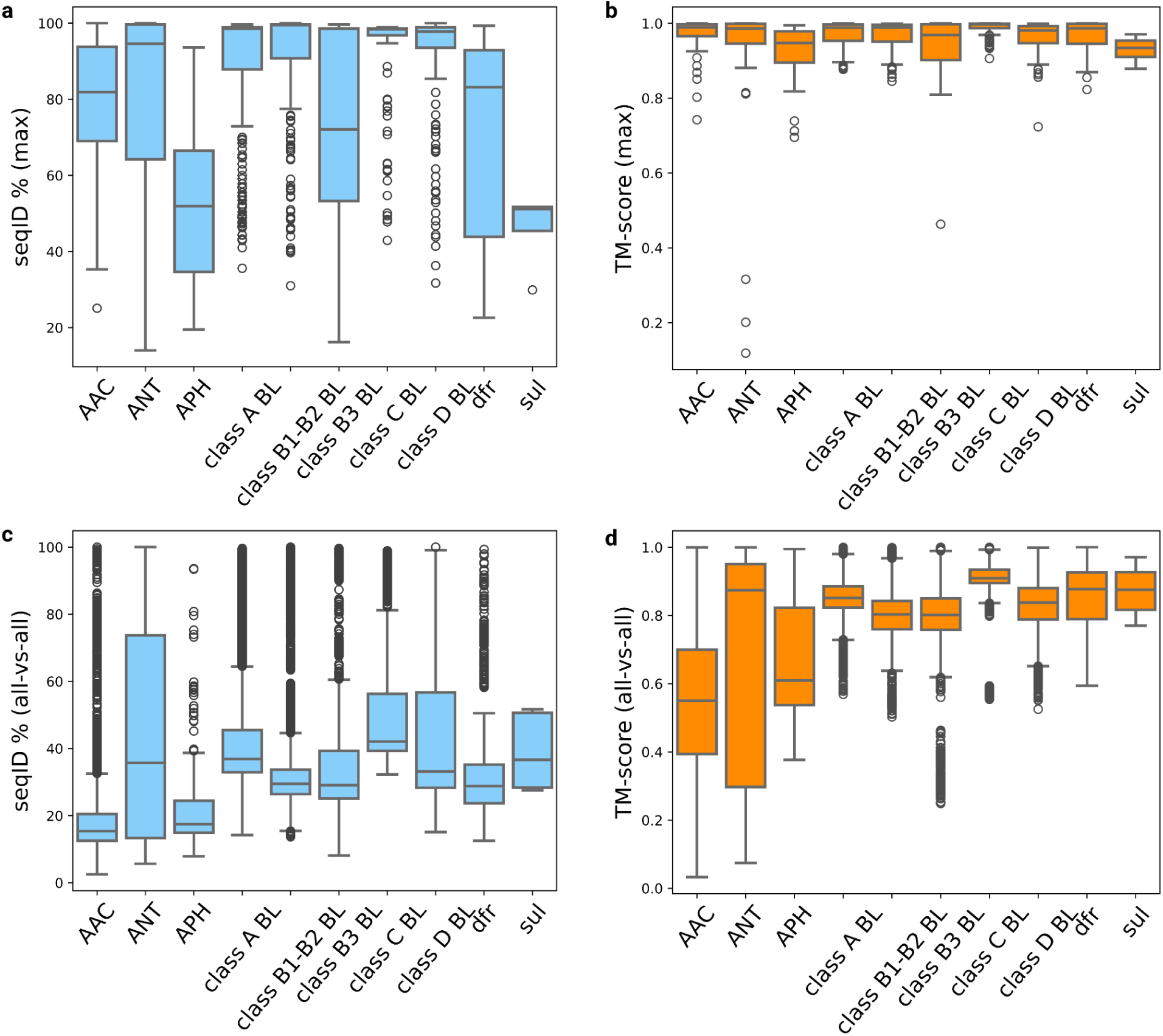
Sequence-structure similarities among antibiotic resistant protein (ARP) structures encoded by well-known ARG classes. **a**, Boxplots of pairwise maximum aa sequence identities (seqIDs) and **b**, their corresponding TM-scores. **c**, Boxplots of pairwise (all-versus-all) seqIDs and **d**, their corresponding TM-scores. ARG classes: aminoglycoside acetyl-transferases (AAC, number of pairwise comparisons (*n*) = 6,228), aminoglycoside nucleotidyl-transferases (ANT, *n* = 2,169), aminoglycoside phospho-transferases (APH, *n* = 1,190), class A β-lactamases (class A BL, *n* = 87,320), class B1/B2 BL (*n* = 41,006), class B3 BL (*n* = 3782), Class C BL (*n* = 27,060), Class D BL (*n* = 48,180), dihydrofolate reductases (dfr, *n* = 2,070), and mobile dihydropteroate synthases (*sul*, *n* = 12). Pairwise seqIDs and TM-scores were calculated using Foldseek under default settings, with TM-scores normalised by the subject length.

Nonetheless, ARP structures typically evolve from existing proteins (proto-resistance genes (Morar & Wright, 2010)) performing primary functions, and are often involved in the target pathway of an antibiotic (Baquero et al., 2021). Proto-resistance genes exhibit high TM-scores to ARP structures, however lack ARG function. Small changes in critical aa residues can make the difference between a true ARG, from a proto-resistance gene which does not confer clinical resistance but could evolve there (for e.g., the evolution of mobile sul genes from chromosomal *folP* (Venkatesan et al., 2023)). Therefore, simply prioritising proteins with high TM-scores to find novel ARP structures would likely result in the generation of a high false positive rate (FPR) (Kaderabkova et al., 2022).

Conversely, ARGs can also have reduced TM-scores to ARP structures, as depicted primarily for AAC’s, ANT’s, APH’s, and class B3 β-lactamases (Figure 1d). This could be explained by ARP structures performing functions by similar localised structural motifs encoded by important aa’s (Koehler Leman et al., 2023). Proteins which contain these motifs without structural conservation in other regions would have reduced TM-scores to ARP structures, so only prioritising proteins with high TM-scores would also result in a high false negative rate (FNR).

By focussing on functionally important protein regions of ARGs, we developed a novel, precise, and computationally efficient method for identifying previously unrecognised ARGs. ARG-PASS (ARG-PAirwise Sequence vs Structure) leverages a one-class support vector machine (SVM) trained on pairwise primary and tertiary distributions of structurally conserved regions of proteins encoded by ARGs. We applied ARG-PASS to six reference strains of the Human Microbiome Project (HMP), and predicted 16 novel ARGs. Of these, 9 were selected for experimental verification in *Escherichia coli* and all had detectable activity: one APH(6’) from *Pseudomonas aeruginosa*; two class B3 β-lactamases from *Burkholderiales* sp. and *Acinetobacter radioresistens*; a cephalosporinase from *Acinetobacter junii*; four *dfr* genes; and a β-lactam-resistant penicillin binding protein (PBP) from *A. radioresistens*. Seven reached MICs at Clinical and Laboratory Standards Institute (CLSI) resistant breakpoints (CLSI, 2024a), and we posit the remainder represent ‘pre-resistance’ genes, with activity below CLSI resistant breakpoints but at risk of becoming clinically relevant in the future. While our primary dataset was HMP reference strains, ARG-PASS also predicted a highly divergent *phnP* gene (metallo-β-lactamase superfamily protein) directly from the AlphaFold database (AFDB), which conferred a clinically relevant MIC against ampicillin when expressed in *E. coli.* The average amino acid (aa) sequence identity (seqID) of functionally confirmed genes to known ARGs is 46%, highlighting the ability of ARG-PASS to uncover genes with low homology to existing ARGs and potentially a large reservoir of previously uncharacterised ARGs with clinical relevance.

## Results

### Development of the ARG-PASS method

Experimental validation remains the main bottleneck in the ARG discovery pipeline. High-precision computational methods are therefore required to identify potential novel ARGs, including those with low sequence identity to known resistance genes. We hypothesised that the resistance function of a potentially novel ARP structure can be inferred by comparing its primary and tertiary structural features to those of structurally conserved regions of known ARP structures. Specifically, we propose that similarity to conserved folds shared across functionally related ARGs, would be a simple and precise method of detecting novel ARGs.

This framework comprises three steps.

1. For each ARP structure within a functionally coherent subgroup (cluster) of an ARG class, extract a consensus structural fold conserved across the subgroup;
2. For each conserved fold, quantify the pairwise (all-versus-all) structural similarity using the TM-score (to assess overall fold similarity), and the seqID at aligned positions (to capture conservation of functionally important residues);
3. A query protein structure is compared against the conserved folds of the ARPs within each cluster using the same metrics. If the query protein structure has any TM-score and seqID values comparable to or exceeding those observed among the cluster, we infer that it also shares the conserved structural features and therefore likely encodes the same resistance function.

We integrated these steps into a single computational framework and validated predictions using standardised expression systems combined with antimicrobial susceptibility testing across CLSI breakpoints. To this end, ARG-PASS (ARG-PAirwise Sequence vs Structure) calculates pairwise primary-sequence and tertiary-structure similarities within conserved, functionally important protein regions (*high*-*lDDT*) of clustered ARPs. ARG-PASS generates a two-dimensional distribution of seqIDs and TM-scores that reveals the sequence–structure landscape of conserved regions of ARG families.

A one-class SVM^8^ was trained to learn the decision boundary surrounding the training distribution. A query ARP (qARP) structure is aligned to each *high-lDDT* ARP structure, and its derived seqID and TM-score are calculated. If any points lie within the decision boundary (DB) of the 𝑥, 𝑦 distribution, it is predicted as functional; if any points lie outside the DB, it is flagged as an outlier, predicting it as non-functional (Figure 2). We validated the approach on an existing protein structure-based method used to identify ARGs from the human intestinal microbiota, called Pairwise Comparative Modelling (PCM) (Ruppé et al., 2019). Among ARG classes with positive and negative functional synthesis results and seqIDs below 80% to known ARGs (AAC(6’), APH, class A and class B β-lactamases, *n* = 33), the precision increased from 0.64 across both ‘fair’ and ‘good’ predictions of PCM, to 0.81 of ARG-PASS, with an accuracy of 0.76 (Table 4, Additional file 1: Table S5). Furthermore, high precision was retained at low seqID, with 13 out of 16 (0.81) true positive predictions correct at seqIDs below 31% to known ARGs (see Methods).

**Figure 2.**
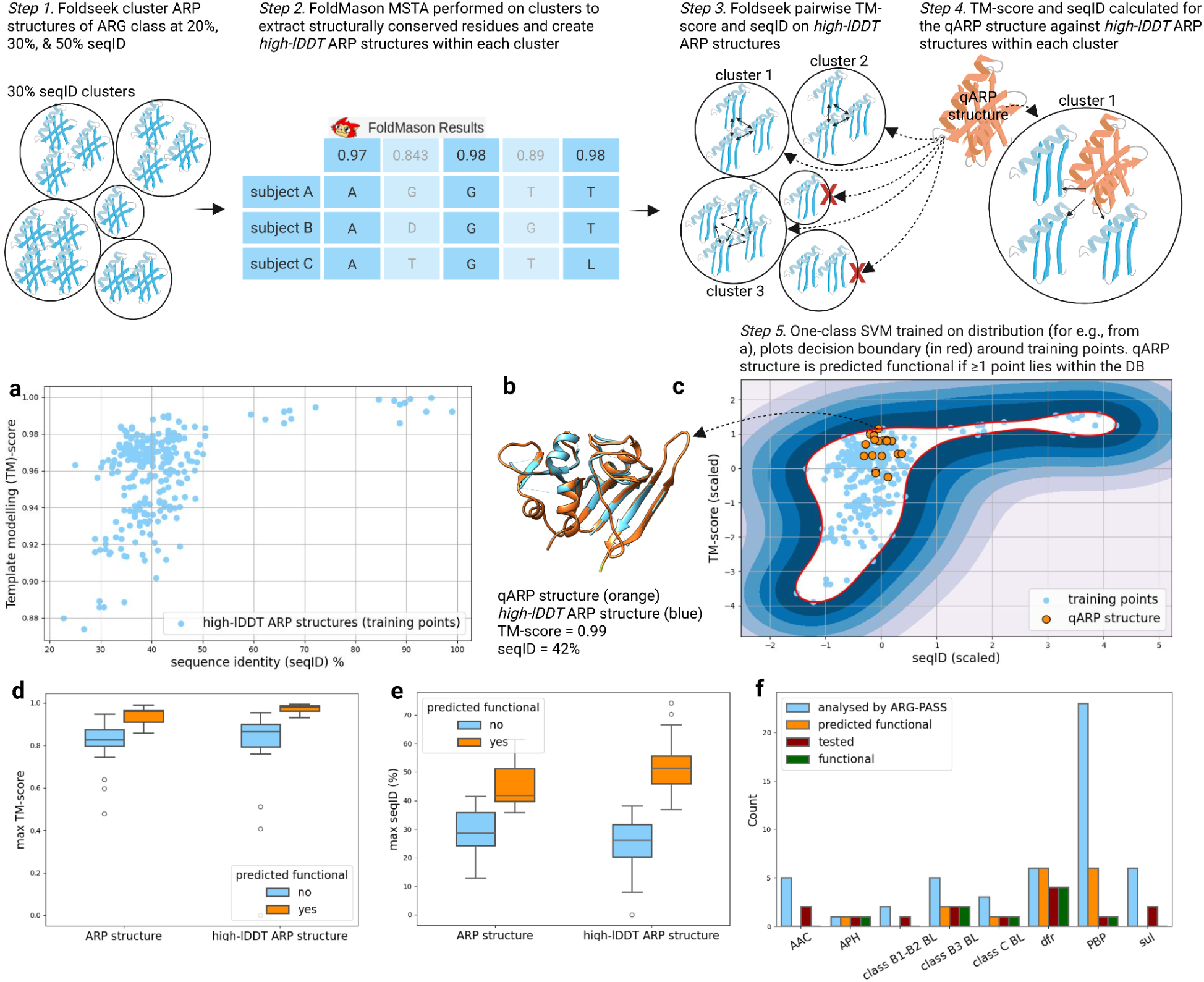
Overview of ARG-PASS. *Step 1*. ARP structures for an ARG class are clustered at 20%, 30%, and 50% seqID using Foldseek. *Step 2.* For each cluster, a MSTA is performed using FoldMason. Aligned residues with a per-column local Difference Distance Test (lDDT) score ≥ Q50, Q75, or Q90 percentiles of the per-column lDDT distribution are extracted to create *high-lDDT* ARP structure .pdb files for each ARP structure. *Step 3.* The pairwise (all-vs-all) seqIDs vs TM-scores are calculated for *high-lDDT* ARP structures within clusters using Foldseek TM-align. *Step 4.* qARP structures are assigned to ARP cluster levels from Step 1 based on their highest seqID to an ARP structure (determined by Foldseek). SeqIDs and TM-scores of the qARP structure to *high-lDDT* ARP structures in each cluster of the level are calculated using Foldseek TM-align. **a**, Example distribution of pairwise seqIDs (𝑥) vs TM-scores (𝑦) for a 30% seqID *high-lDDT* ARP structure cluster of *dfr’s*. *Step 5* (**c**). A one-class SVM is trained on the 𝑥, 𝑦 distributions from *Step 3,* the decision boundary (DB, in red) surrounds all training points. The seqIDs vs TM-scores of the aligned region of a single qARP structure against each *high-lDDT* ARP structure from *Step 4* are plotted (data points in orange). If any data points lie on or within the DB, the qARP is classed as functional (**b**, example), otherwise the qARP is classed as non-functional. **d**, **e**, Highest TM-score and seqID between qARP structures and ARP structures vs *high-lDDT* ARP structures after ARG-PASS (Mann-Whitney test between yes/no predictions of each condition: *p* < 0.001). **f**, Numbers of qARPs from HMP reference strains analysed in this study. For further details see Methods. Figure created with BioRender.com.

### ARG-PASS identified novel ARGs from Human Microbiome Project reference strains

Protein structures encoded by query antibiotic resistant proteins (qARPs) originating from six HMP reference strains (The Human Microbiome Jumpstart Reference Strains Consortium et al., 2010) (Additional file 1: Table S1), were downloaded directly from the AFDB if available or computed by AlphaFold2 (AF2) or AlphaFold3 (AF3), representing qARP structures. ARG-PASS then functionally predicted qARP structures using seqID, TM-score distributions of *high-lDDT* ARP structures (one-class SVM models, see Methods) created for well-known ARG families (aminoglycoside acetyl-transferases (AAC), aminoglycoside nucleotidyl-transferases (ANT), aminoglycoside phospho-transferases (APH), class A, B, C, and D β-lactamases, dihydrofolate reductases (*dfr*), class 1, 2, and 3 β-lactam-resistant penicillin binding proteins (PBP’s), and mobile sulfonamide-resistant dihydropteroate synthetases (*sul*)).

In total, 16 qARPs were classed as functional by ARG-PASS, predicted to provide resistance to streptomycin, β-lactams, and trimethoprim (Figure 2f, Additional file 1: Table S2). The average highest TM-score (0.97) and seqID (54%) of qARP structures predicted as functional shared with *high-lDDT* ARP structures, increased from comparison to full-length ARP structures (TM-score of 0.94 and seqID of 45.7%).

Conversely, the average highest TM-score of qARP structures predicted as non-functional to *high-lDDT* ARP structures from ARP structures remained consistent (0.82) and seqIDs actually decreased (29.7% to 25.5%), perhaps underscoring the ability of the algorithm to distinguish from qARPs which don’t have functionally important primary and tertiary structural features (Figure 2d and 2e).

Nine qARPs predicted as functional by ARG-PASS and belonging to five ARG classes (APH, *dfr*, class B3 and C β-lactamases, and β-lactam-resistant PBP), 5 qARPs belonging to ARG classes assessed by ARG-PASS without any predicted as functional (Figure 2f, class B1/B2 β-lactamases, AAC, and *sul*), and positive controls (not already available in the Minimal Antibiotic Resistance Platform), were synthesised and cloned into appropriate plasmids (pGDP1, pGDP2, or pGDP3) and expressed in *Escherichia coli* BW25113 (Δ*bam*BΔ*tolC*) for functional verification (Cox et al., 2017). We used the efflux deficient *E. coli* strain for expression of qARPs, so any activity could not be attributed to efflux pumps. Activity was detected in all qARPs predicted as functional against their respective antibiotics (Additional file 1: Table S2), with minimum inhibitory concentrations (MICs) determined in liquid media, by broth microdilution (BM) and solid media, by a spot plating assay (SPA) (Table 1). Genomic context analysis suggests qARPs with detected activity represent stable chromosomally encoded components of gut microbiome genomes rather than within MGEs. No activity was detected in qARPs predicted as non-functional by ARG-PASS (Additional file 1: Table S2, MIC data not shown).

**Table 1.**
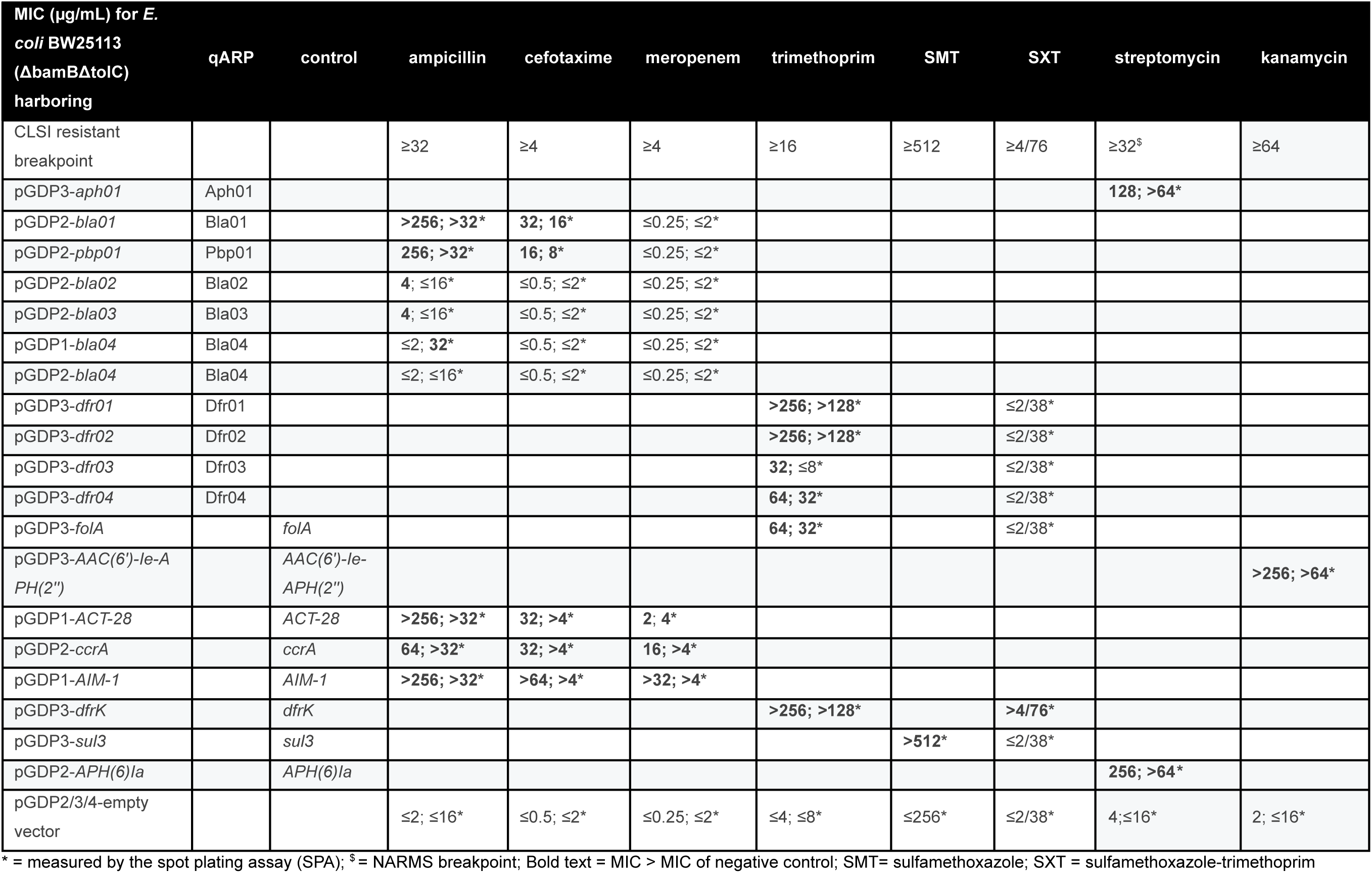
Functional confirmation results of qARPs and controls. All MICs were determined by Broth Microdilution (BM) unless marked.

### qARPs conferred MICs at CLSI resistant breakpoints

#### β-lactamases

For ampicillin, the negative control strain exhibited an MIC of 2 µg/mL. According to CLSI guidelines, susceptibility is defined as MIC ≤8 µg/mL and resistance as MIC ≥32 µg/mL. Expression of *bla01* increased the MIC to >256 µg/mL measured by BM and to >32 µg/mL measured by the SPA, representing up to a 128-fold increase relative to the negative control, and places the strain in the resistant category for ampicillin (Table 1). For cefotaxime, the negative control strain exhibited an MIC of ≤2 µg/mL, while the CLSI susceptibility and resistance breakpoints are ≤1 µg/mL and ≥4 µg/mL, respectively.

Expression of *bla01* increased the cefotaxime MIC to 32 µg/mL by BM and 16 µg/mL by the SPA, also placing the strain in the resistant category for cefotaxime (Table 1).

Bla01 is a potential cephalosporinase from *Acinetobacter junii* SH205 with 64.2% seqID to ADC-8 (Additional file 1: Table S3). ARG-PASS predicted the highest TM-score (0.99) and seqID (73.8%) of Bla01 is shared with RSC1-1_Q50_lddt_ (Figure 3a and Figure 3b).

**Figure 3.**
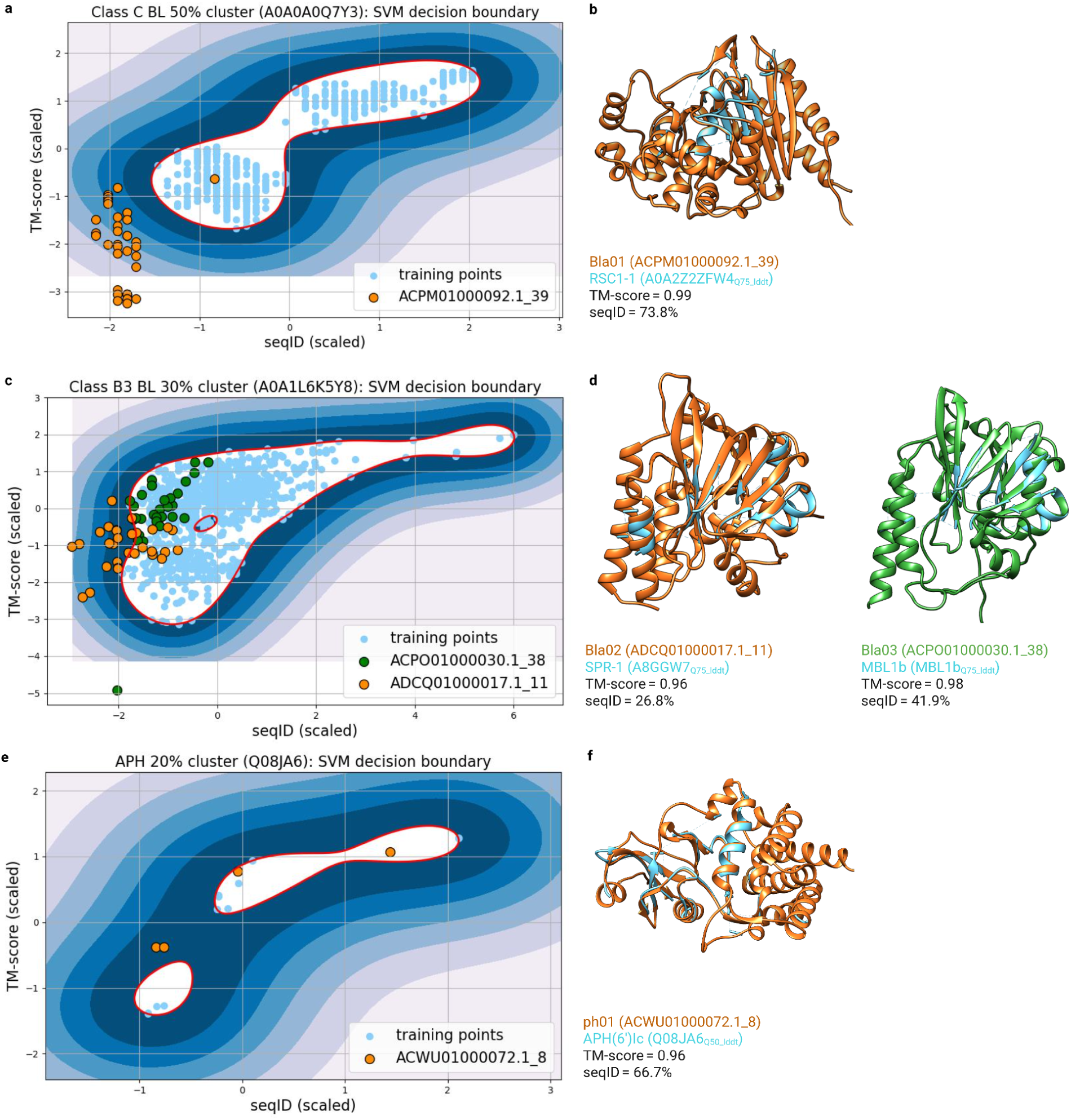
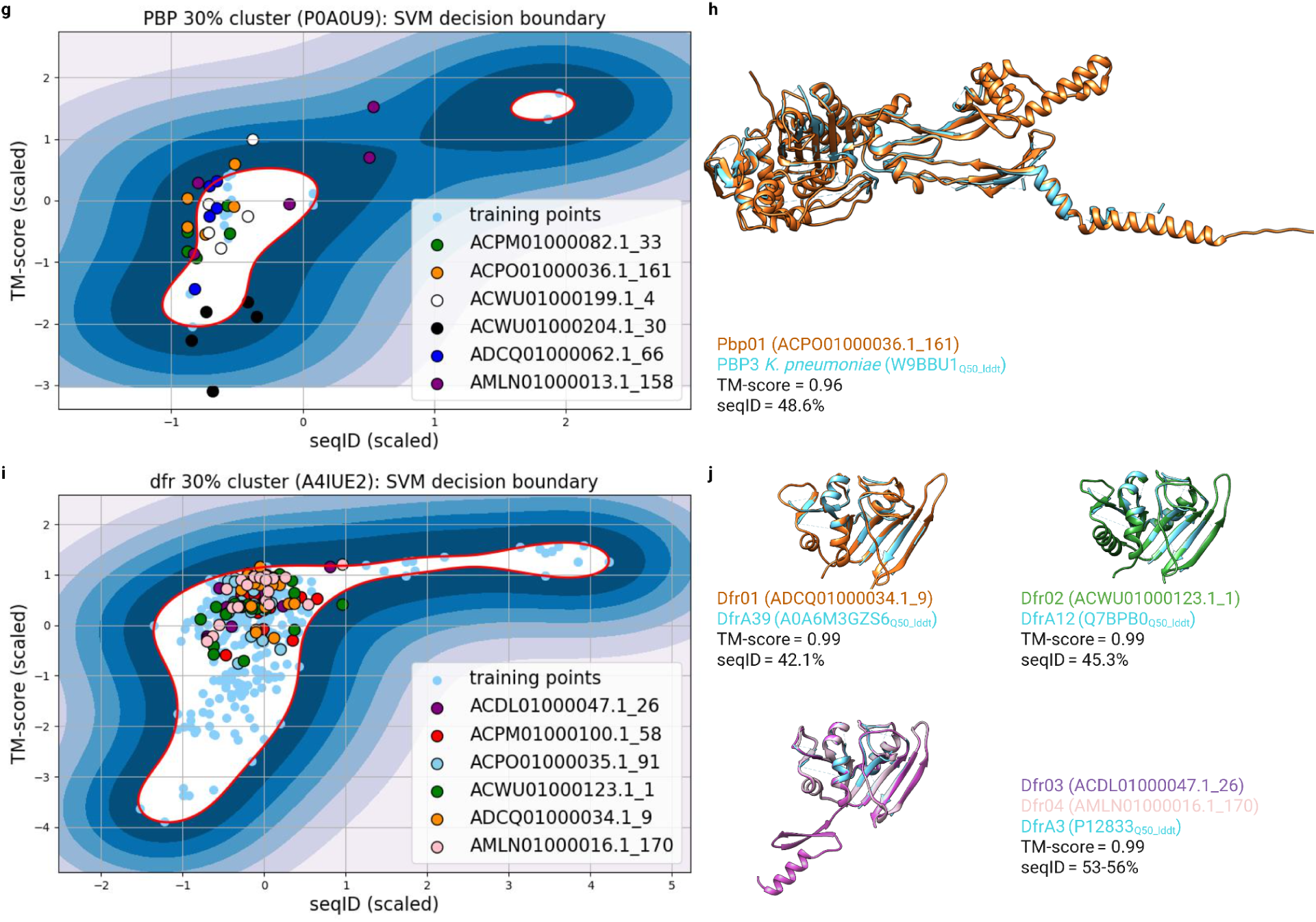
**a, c, e, h, i**, One-class SVM models for qARP structures predicted as functional by ARG-PASS, each depicted with at least one data point within the DB (in red) surrounding the training distribution. **b, d, f, h, j**, qARP structures with activity following functional verification are overlaid with the *high-lDDT* ARP structure with the highest TM-score. Protein sequences of qARPs and fasta files of Foldseek structural alignments of qARP structures and *high-lDDT* ARP structures with the highest TM-score are available in Additional file 2 and 3. One-class SVM plots were made using scikit-learn (Pedregosa et al., 2011) and AlphaFold structures visualised with UCSF Chimera (Meng et al., 2006).

RSC1-1 was identified from Indian river sediments (Marathe et al., 2018). Based on low sequence homology to existing cephalosporinases and elevated MICs against β-lactams, we tentatively propose a name for Bla01 of AJDC-1. Catalytic motifs known to be responsible for β-lactam hydrolysis are conserved in AJDC-1 (^64^SXSK, ^150^YXN, and ^315^KTG) (Philippon et al., 2022) and were captured as conserved regions of *high-lDDT* ARP structures in clusters of known cephalosporinases by ARG-PASS (Figure 3b, Additional file 3: Alignment 1).

*Acinetobacter* sp. are common sources of *ampC* cephalosporinases, in particular ADC (*Acinetobacter*-derived cephalosporinases) variants which often originate in *A. baumannii* (Hujer et al., 2005). AJDC-1 represents the first cephalosporinase functionally confirmed from HMP reference strain *A. junii* SH205, which may be the naturally occurring cephalosporinase from this species. Strains of *A. junii* are not commonly associated with disease, but it has been reported as an emerging One-Health pathogen (Aguilar-Vera et al., 2024). Furthermore, the earliest observations of *bla*_NDM-1_ and *tet(X3)* originate from *A. junii* (Grenier et al., 2024a; 2024b).

Phylogenetic inference of AJDC-1 with ADCs and other closest related cephalosporinases from *Acinetobacter* sp. (including those without MIC data obtained from the Beta-Lactamase DataBase (BLDB) (Naas et al., 2017) and NCBI), confirms its phylogenetic proximity to ADC-8, functionally confirmed in 2007 from *Acinetobacter baylyi*, however is closest to the potential AmpC cephalosporinase from *A. beijerinckii* (not present in CARD). Both AJDC-1 and ADC-8 are divergent from other functionally confirmed cephalosporinases (Zhao & Hu, 2012) (Additional file 4: Figure S1).

#### Streptomycin resistance

The negative control strain exhibited an MIC of 4 µg/mL against streptomycin as measured by BM. There are currently no CLSI breakpoints established for streptomycin, however the National Antibiotic Resistance Monitoring Network for Enteric Bacteria (NARMS) record an interpretative susceptible MIC of ≤8 µg/mL and resistant MIC of ≥32 µg/mL for years tested between 2014-2019 (CDC, 2025). Expression of *aph01*, a potential APH(6’) from HMP reference strain *P. aeruginosa* 2_1_26, increased the streptomycin MIC to 128 µg/mL by BM and MIC >64 µg/mL by the SPA, similar to the positive control, APH(6’)Ia.

Aph01 shares the highest seqID of 52.6% with APH(6)-Ic (Additional file 1: Table S3) and ARG-PASS identified a TM-score of 0.96 and seqID of 66.7% of Aph01 to APH(6)1c_Q50_lddt_ (Figure 3e and 3f). Four known streptomycin-resistant APH(6’)’s are listed in CARD and all clustered at 20% seqID in the ARG-PASS pipeline (Figure 3e). Along with APH(6’)Ic and APH(6’)Id, Aph01 likely represents the third known from *P. aeruginosa,* and fifth known APH(6’) gene. As such, we tentatively designate its name as APH(6’)Ie. All three APH(6’) enzymes identified from *P. aeruginosa* (including APH(6’)Ie presented here) are phylogenetically closer than APH(6’)Ia and APH(6’)lb, from *Streptomyces griseus* and *S. glaucescens*, respectively (Additional file 4: Figure S2).

#### Trimethoprim and trimethoprim-sulfamethoxazole resistance

The CLSI defines susceptibility and resistance to trimethoprim (TMP) as MIC’s of ≤8 µg/mL and ≥16 µg/mL, respectively. When expressed in *E. coli*, *dfr01, dfr02* and *dfr04*, all encoding potential TMP-resistant dihydrofolate reductases (DHFRs) from *Burkholderiales* sp. 1_1_47, *P. aeruginosa* 2_1_26, and *K. pneumoniae* WGLW3, respectively, conferred clinical resistance to TMP by BM (MIC >256 µg/mL for Dfr01 and Dfr02, and MIC 64 µg/mL for Dfr04) and by the SPA (MIC >32 µg/mL for Dfr01 and Dfr02, and MIC 32 µg/mL for Dfr04). These MICs were similar to the positive control, *dfrK*, and up to 64-fold higher than the negative control (MIC ≤4 µg/mL by BM) (Table 1). Dfr03 from *E. coli* 53FAA also conferred elevated MICs to TMP, however only when measured by BM. Differential growth dynamics in liquid versus solid media potentially influenced the observed activity of Dfr03 and has been reported previously for the susceptibility of *Campylobacter* sp. to sulfamethoxazole-trimethoprim (SXT) (Luber et al., 2003). ARG-PASS determined Dfr01 and Dfr02 have highest TM-scores of ∼0.99 to DfrA39_Q50_lddt_ and DfrA12_Q50_lddt_ , respectively, and Dfr03 and Dfr04 share their highest TM-score of ∼0.99 with seqID of 53-56% to DfrA3_Q50_lddt_ (Figure 3j, Additional file 1: Table S2).

The resistant category to SXT is defined by the CLSI as an MIC of ≥4/76 µg/mL. The strain harbouring positive control *dfrK* also conferred clinical resistance to SXT. As a structural analogue of p-aminobenzoic acid, sulfamethoxazole competitively inhibits dihydropteroate synthetase (DHPS), which is an earlier step than where TMP acts in the folate synthesis pathway (Ovung & Bhattacharyya, 2021). Clinical resistance to SXT is commonly reported by both a *dfr* and *sul* gene (*sul1*-*sul4*) and less likely by the presence of only either (Shin et al., 2015, Poey et al., 2024). Interestingly, a clinical MIC to SXT was not observed for the strain harbouring a known mobile *sul* gene (*sul3)*, as opposed to *dfrK,* which may have clinical relevance (Table 1).

We investigated whether elevated TMP MICs were caused by over production DHFR which can titrate out TMP due to its mechanism as a competitive inhibitor, particularly under the control of the *bla* promoter in the pGDP3 plasmid (Palmer & Kishony, 2014; Soo et al., 2011). We cloned and expressed the wild-type (WT) *E. coli* chromosomal *Ec*DHFR (encoded by *folA*) into pGDP3. *Ec*DHFR conferred a similar TMP MIC to *dfr04* by both measurement protocols, and conferred a higher MIC than *dfr03*. *Ec*DHFR actually shares 100% seqID with Dfr03, apart from an additional 37 residues constituting an α-helix at the N-terminus in Dfr03 (Figure 3j). This may have played a role in restricting the expression of *dfr03* and thus the lower MICs recorded compared to *Ec*DHFR. Furthermore, Dfr04 shares a seqID of 92% to Dfr03. After sequence alignment, substitutions present in Dfr04 from Dfr03 are V41I, H45L, I61V, L62I, Q65K,T68S, T73Q, K76S, D79E, P89E, E129D, D132E, and H149N. Further research is needed to deduce if these substitutions affected the variable TMP MICs observed.

We searched 10 open reading frames (ORFs) up and downstream of genes encoding Dfr01-Dfr04 for signatures of MGEs, which had at least 80% seqID (over 90% query and subject length), to proteins within the MobileOG-db database (Brown et al., 2022). The absence of exclusively MGE-associated sequences prevented confident inference of mobility for Dfr01-Dfr04. Moreover, contigs containing *dfr01*-*dfr04* from their respective hosts are predicted to originate from chromosomal DNA by PlasmidHunter (Tian et al., 2024). This suggests Dfr01-Dfr04 are DHFR products of chromosomally encoded *folA* genes in their respective HMP reference strains.

The strains expressing *dfr01* and *dfr02* exhibited three-fold higher TMP MICs than *dfr03*, *dfr04*, and *Ec*DHFR, and conferred comparable MICs to the positive control *dfrK*. This phenotype is consistent with intrinsic enzyme properties such as reduced TMP affinity (higher K_i_) and/or increased catalytic efficiency (k_cat_) relative to Dfr03, Dfr04, and *Ec*DHFR. Steady-state kinetic assays would be required to confirm these mechanisms. When Dfr01-Dfr04 are structurally aligned using FoldMason with the remaining Dfr’s in their one-class SVM cluster for which they were predicted functional (Additional file 3: Alignment 7), no known resistance-conferring mutations are present across both queries and subjects (Manna et al., 2021; Tamer et al., 2019). This provides support for the common mechanism of TMP-resistance in known mobile *dfr* genes: expression of decontextualised *folA*. It is also unsurprising that ARG-PASS predicts Dfr01-Dfr04 as functional, given it was trained on decontextualised *folA* genes (i.e., Dfr proteins from the CARD). There is alarming potential for these enzymes to confer clinical resistance to TMP when expressed in a low-copy plasmid by a promoter of similar strength to the *bla* (as observed under these experimental conditions), and may provide evidence for the ubiquity of mobile Dfr’s currently known.

Phylogenetic analysis with the remaining Dfr proteins in the cluster for which Dfr01-Dfr04 were predicted functional, indicates they are distinct from known TMP-resistant Dfr’*s* (Additional file 4: Figure S3). Dfr02 from *P. aeruginosa* is sister to Dfr03 and Dfr04, and is closest related to the integron-encoded DfrA3 from *E. coli* (Brolund et al., 2010). Strains of *P. aeruginosa* are often intrinsically resistant to TMP by low outer membrane permeability, efflux pumps, and the integron-encoded *dfrB5* (Faltyn et al., 2019; Pang et al., 2019). Being the most divergent identified in this study, Dfr01 is outgrouped from Dfr02-Dfr04 (Additional file 1: Table S3; Additional file 4: Figure S3).

*Burkholderiales* sp. may represent a source of *dfrB* genes and to the best of our knowledge, Dfr01 represents the first report of *folA* from *Burkholderiales* sp. conferring clinical resistance to TMP when expressed in *E. coli* (Kneis et al., 2024).

#### A PBP3 homolog from *A. radioresistens* was functionally confirmed to provide resistance to β-lactams

The strain harbouring Pbp01, a predicted penicillin binding protein (PBP) from *A. radioresistens* SH164, conferred MICs above CLSI resistant breakpoints against ampicillin by BM (MIC 256 µg/mL) and the SPA (MIC >32 µg/mL), and cefotaxime by BM (MIC 16 µg/mL) and the SPA (MIC 8 µg/mL, Table 1). Based on results of the Diamond search against the custom CARD database, Pbp01 (ACPO01000036.1_161) shares a seqID of 40.7% to the WT PBP3 from *Klebsiella pneumoniae* (Additional file 1: Table S3). Computational prediction of membrane-associated proteins have lower relative confidence (Agarwal & McShan, 2024) and PBP’s are longer than most ARPs (i.e., >500 aa’s). Foldseek (used in the ARG-PASS pipeline) structurally aligned distant residues of Pbp01 to the *high-lDDT* ARP structure of PBP3 from *K. pneumoniae* (AlphaFold: W9BBU1) with a TM-score of 0.96 and seqID of 48.6% (Figure 3h, Additional file 1: Table S2). To the best of our knowledge, Pbp01 represents the first functionally verified report of a PBP3 from *A. radioresistens* conferring β-lactam resistance when expressed in *E. coli*.

Point mutations in PBPs reduce the affinity of β-lactam antibiotics for their targets (Zapun et al., 2008). ARP structures used to create one-class SVM models for PBPs, were encoded by the representative WT sequences of PBP1–PBP3 in the protein-variant model in CARD, without introducing resistance-conferring mutations.

This approach was considered appropriate because structural alignment of PBPs containing large numbers of diverse point mutations, often with highly localised effects, would be unlikely to yield a reliable consensus of structurally conserved regions (*high-lDDT*). For example, CARD lists 30 resistance-conferring point mutations/insertions for *Helicobacter pylori* PBP1 mutants (ARO: 3007060). The experimental confirmation of a predicted resistant PBP3 homologue, Pbp01, using a TM-score vs seqID distribution of conserved structural regions of WT PBPs, suggests these regions possess a consensus structural backbone, which, in addition to point mutations, could be important for the function of β-lactam resistant PBPs. Five other PBPs were predicted as functional in this study, originating from *A. junii*, *P. aeruginosa*., *K. pneumoniae* subsp. *pneumoniae*, and *Burkholderiales*. sp., with TM-scores over 0.93 and seqIDs over 45% to known PBP3’s. It is difficult to assert if these likely represent true β-lactam-resistant PBP3’s given only one PBP3 was experimentally tested; however, further research is warranted into the β-lactam susceptibility of these strains.

Reports linking PBPs to β-lactam resistance need further elucidation in *Acinetobacter* sp. (Sethuvel et al., 2023; Zapun et al., 2008). Further research is needed to deduce if the observed phenotype in Pbp01 is derived from point mutations and/or a reduced affinity for β-lactams (Zapun et al., 2008). OXA-905, only known from *P. aeruginosa* in CARD, was also identified in *A. radioresistens* in this study (Additional file 1: Table S3) suggesting this strain could already be resistant to penicillins and/or cephalosporins (Evans & Amyes, 2014). A Blastp search against the NCBI database reveals Pbp01 shares the highest seqID of 85% to a PBP3 from the clinical pathogen *A. baumannii* (NCBI: MFY0112749.1). We could not ascertain whether this PBP3 from *A. baumannii* also provides resistance to β-lactams. Their significant homology shared poses a risk for *pbp01* to horizontally transfer to *A. baumannii*, as occurred with the *mecA* gene (methicillin resistant PBP2) to *Staphylococcus aureus* (Utsui & Yokota, 1985).

#### Pre-resistance genes encode qARP’s with activity but MICs below CLSI resistant breakpoints

*E. coli* strains expressing *bla02* and *bla03*, encoding potential metallo-β-lactamases (MBLs) from *Burkholderiales* sp. 1_1_47 and *A. radioresistens,* respectively, conferred a two-fold increase in the ampicillin MIC determined by BM (MIC 4 µg/mL) against the negative control (MIC ≤2 µg/mL). ARG-PASS determined both Bla02 and Bla03 share the highest structural similarity in conserved protein regions to class B3 MBLs, SPR-1_Q75_lddt_ and MBL1b_Q75_lddt_, respectively (Figure 3d, Additional file 1: Table S2).

Whilst these qARPs had activity against ampicillin compared to the negative control, they did not confer an MIC at the CLSI resistant breakpoint, and within the susceptible category with an MIC ≤8 µg/mL. We term these ‘pre-resistance’ genes – genes which may have high potential to evolve into ARPs (i.e., with clinical relevance). They are likely further along the evolutionary path to clinical resistance than proto-resistance genes, and we posit they constitute another subset of the resistome (Morar et al., 2009; Morar & Wright, 2010). Moreover, we hypothesise pre-resistance genes like Bla02 and Bla03, may occupy a critical area within the fitness landscape of ARGs, where they are maintained in bacterial populations with relatively low fitness cost (for e.g., compared to canonical MBLs (López et al., 2019a)), under commonly encountered, subclinical exposure to antibiotics (Figure 4b).

**Figure 4.**
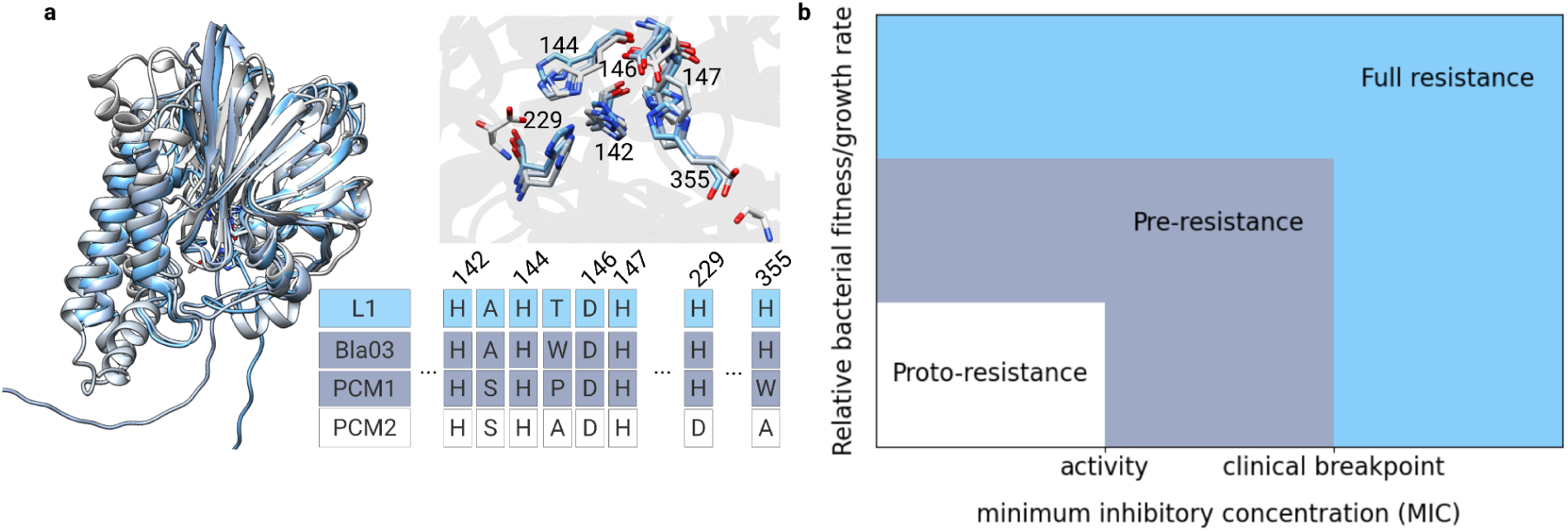
Conceptual Fitness-MIC landscape of the resistome as determined by Zn active site architecture of class B3 MBL’s. **a,** L1, Bla03, PCM1, and PCM2 exhibit overall fold similarity however PCM1 and PCM2 lack critical Zn- binding histidine residues (full Chimera structural alignment is available in Additional file 3: Alignment 11). **b**, Fitness-MIC landscape comprising components of the Resistome within a continuum, ranging from proto-resistance elements with no activity (for e.g., PCM2), through pre-resistance genes that confer detectable MIC increases yet without clinical relevance (for e.g., Bla03 and PCM1), to fully evolved resistance determinants (for e.g., L1).

It should be noted an efflux-pump deficient *E. coli* BW25113 (Δ*bam*BΔ*tolC*) was used to functionally verify qARPs to ensure all elevated MICs are attributed directly to the query enzyme. Although Bla02 and Bla03 identified in this study alone are not sufficient to confer clinically relevant MICs, they may act in concert with other molecular determinants when selected by higher antibiotic dosing regimes and subsequently reach CLSI resistant breakpoints, for example, alongside increased efflux of carbapenems (Pai et al., 2001; Tewawong et al., 2025). In addition to potential catalytic inefficiency compared to other MBLs, the amount of functional perisplasmic enzyme may have also negatively impacted the observed phenotype (Socha et al., 2019) and can be influenced by an incompatible signal peptide with the expression system (López et al., 2019). This is also supported by the Zn^2+^-binding active site encoded by residues H116, H118, D120, H121, H196, and H263 in class B3 MBLs (Boyd et al., 2020), conserved in Bla02 and Bla03 (Figure 4a, Additional file 3: Alignment 13). As such, higher MICs may be observed in the native host of the gene.

#### Structural alignment suggests a PCM-confirmed class B3 MBL is a candidate pre-resistance gene

After structural alignment of a functional class B3 MBL designated PCM1 (query MC3.MG291.AS1.GP1.C1053.G2 in Ruppé et al., 2019) and a non-functional class B3 MBL designated PCM2 (protein MC3.MG13.AS1.GP1.C10091.G28 in Ruppé et al., 2019), with the known class B3 MBL, L1 and Bla03, reveals that PCM1 lacks histidine residue H263. A tryptophan residue occupies this position (equivalent to H355 in the alignment), and points to a structurally intermediate, class B3 MBL-like architecture (Figure 4a). We therefore speculate that PCM1 may represent a pre-resistance gene rather than a bona fide B3 MBL, a distinction not experimentally verifiable by nitrocefin testing alone (as performed in the PCM study). Moreover, PCM2 lacks two histidine residues: H196 and H263 (equivalent to H229 and H355 in the alignment). Although PCM2 is structurally similar to L1 (Figure 4a), the absence of both residues and no detected activity suggests it represents a proto-resistance gene. Together with catalytic inefficiency and/or potentially incompatible signal peptides noted previously for Bla02 and Bla03, these examples provide putative molecular mechanisms delineating the boundaries between proto-resistance, pre-resistance, and resistance among B3 enzymes. MIC testing of PCM1 against CLSI breakpoints would nonetheless be required to support these hypotheses.

Structural alignment of the putative class B1 MBLs identified by PCM (Ruppé et al., 2019) with canonical class B1 MBLs, NDM-1, VIM-1, and IMP-1, reveals an absence of C221 (critical for Zn^2+^ coordination) across all confirmed proteins, with all other zinc-binding residues conserved (Additional file 3: Alignment 12). After structural alignment to known class B3 MBLs and Bla02 and Bla03, these proteins also possess H121 at the equivalent Zn^2+^ coordination site, which also suggests B3-like active site architecture (Additional file 3: Alignment 13). Notably, they were also predicted as functional class B1 MBLs by ARG-PASS. Although the use of the FoldMason lDDT to threshold conserved structural regions of proteins was sufficient to identify most functional MBLs (including potential pre-resistance genes) from both the PCM study and this paper with high precision, it may lack the resolution to discriminate MBL subclasses at significant levels of sequence divergence and is a limitation planned to be addressed in future work.

### Relaxing ARG-PASS thresholds predicted an evolutionarily divergent MBL-fold protein with ampicillin activity

We next sought to further optimise functional predictions of qARPs by ARG-PASS by skipping the initial Diamond screening of a DNA database against the created database of ARG sequences from the CARD. This was achieved by analysing protein structures directly from the AFDB. We also sought to test the capability of ARG-PASS at predicting proteins with less than 20% seqID to known ARGs and prioritised MBLs given their importance in conferring resistance to last-resort carbapenems. Finally, we prioritised annotations originating from *E. coli* to maximise chances of successful expression in our expression system *E. coli* BW25113 (Δ*bamB*Δ*tolC*). The UniProtKB database (release 2025_02) was queried with the following filter: (taxonomy_id:562) AND (protein_name: Metallo-beta-lactamase). After discarding qARPs with >20% seqID to known MBLs and redundant structures, 50 qARPs were analysed by ARG-PASS. Seven qARPs were predicted as functional class B1/B2 MBLs, however no qARPs were predicted as functional class B3 MBLs. Given class B3 MBLs are the most sequence-divergent subclass of class B MBLs (Krco et al., 2023), this might reflect a limitation of homology-based clustering at a high level of divergence. We therefore conducted an additional analysis excluding prior clustering before obtaining *high-lDDT* ARP structures for class B3 MBLs (see Methods).

Subsequently, 28 qARPs were predicted as functional class B3 MBLs by ARG-PASS, sharing an average highest TM-score of 0.8 and seqID of 17.3% with *high-lDDT* class B3 MBLs. Seven of these were the predicted functional class B1/B2 MBLs, sharing an average highest TM-score of 0.85 and seqID of 22.9% with *high-lDDT* class B1/B2 MBLs (Additional file 1: Table S4).

We selected one qARP(AF/UniProt accession: A0A377FG49), designated Bla04 here, for functional verification. This qARP was one of 13 phosphonate metabolism gene annotations (*phnP)* in the AFDB (carbon-phosphorus lyase complex accessory protein (MBL superfamily)), and β-lactam activity has not been associated before with this gene (Additional file 1: Table S4). No activity compared to the negative control was initially observed with the weaker *lac* promoter (pGDP2), so *bla04* was expressed by the stronger *bla* promoter (pGDP1). In this context, Bla04 conferred an increased MIC against ampicillin when measured by the SPA (MIC 32 µg/mL) compared to the negative control (MIC ≤16 µg/mL), with growth observed at the CLSI intermediate breakpoint of 16 µg/mL. However, an increased ampicillin MIC was not observed compared to the negative control when measured by BM (Table 1).

Bla04 is known from *E. coli* strains 412057 and NCTC9706 (non-HMP reference strains) in UniProt and the *phn* operon in *E. coli* breaks down phosphonates when phosphate is scarce (Yakovleva et al., 1998). After structural alignment using Foldseek of Bla04 to known MBLs, Bla04 shares the highest seqID of 17.4% with SFB-1, which makes it highly divergent from known MBLs.

ARG-PASS revealed Bla04 shares the highest TM-score of 0.8 with a seqID of 19.3% to PNGM-1_Q75-lddt._ (Figure 5a, Additional file 1: Table S4). PNGM-1 was isolated from a deep-sea sediment and has a similar ampicillin MIC of 16 µg/mL when expressed in *E. coli* and determined using MH agar (Park et al., 2018). Only sites H116 and H121 were captured as *high-lDDT* residues in PNGM-1_Q75-lddt_. Noting this dataset was not previously clustered, this likely reflects structural variability in the active site across all class B3 MBLs, and flexible zinc coordination can correspond to different substrate profiles (Krco et al., 2023). Sites H116 and H121 structurally aligned to the active site present in Bla04 by Foldseek in the ARG-PASS pipeline (encoded by residues H76, H80, Figure 5a).

**Figure 5.**
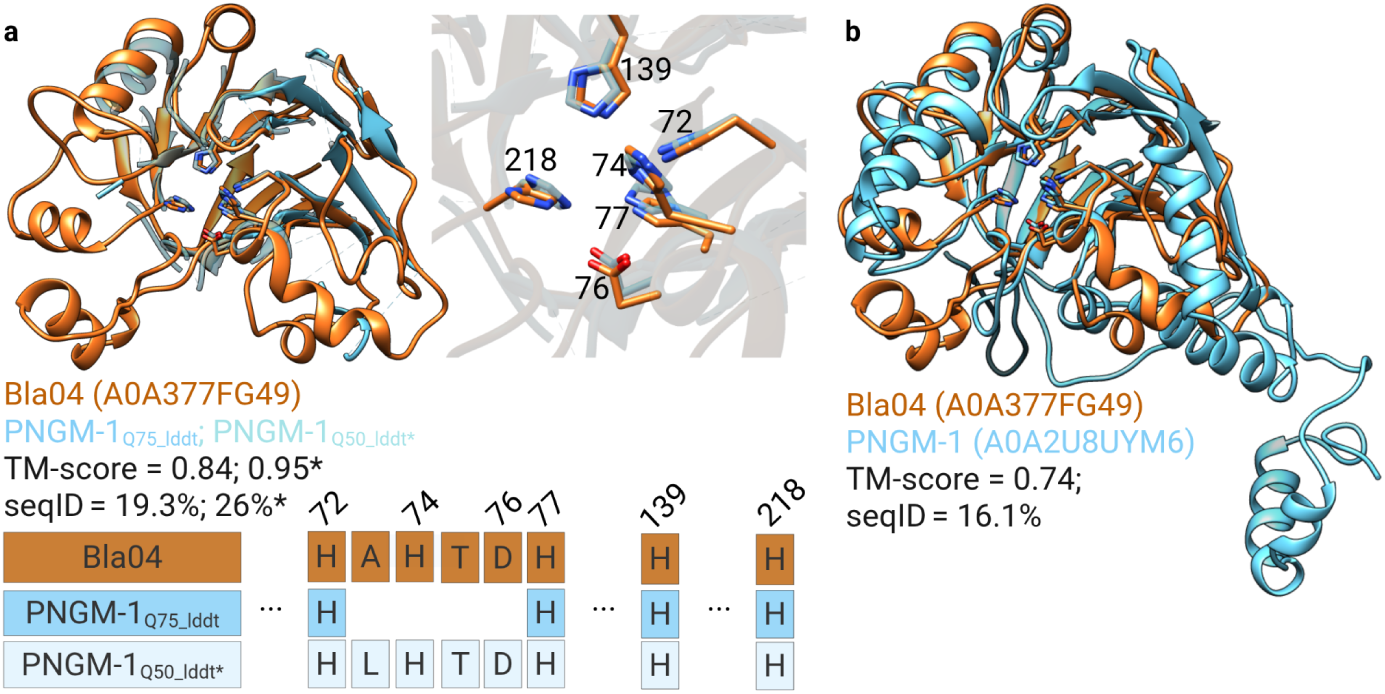
**a**, The Foldseek TM-align alignment (used in ARG-PASS) of Bla04 and the *high-lDDT* ARP structure, PNGM-1_Q75_lddt_ (TM-score of 0.84 and seqID of 19.3%) and to PNGM-1_Q50_lddt*_ (TM-score of 0.95 and seqID of 26%). Aligned active site on Bla04 and PNGM-1_Q50_lddt*_ are displayed. * = *high-lDDT* residues were extracted after pairwise alignment between Bla04 and PNGM-1. **b**, Bla04 aligned to the full length ARP structure of PNGM-1 (TM-score of 0.74 and seqID of 16.1%). Foldseek structural alignments are presented in Additional file 3: Alignment 14 and 15.

When Bla04 and the full-length ARP structure of PNGM-1 are aligned using Foldseek, they share a TM-score of 0.74 and seqID of 16.1% (Figure 5b). After pairwise alignment of Bla04 and PNGM-1 using FoldMason, the Zn^2+^-binding active site in class B3 β-lactamases is conserved in Bla04 (Additional file 3: Alignment 16). To determine if it is potentially contained within a conserved fold relative to PNGM-1, after this pairwise alignment, *high-lDDT* residues of PNGM-1 were extracted to create a new structure file, designated PNGM-1_Q50_lddt*_. Here, the TM-score of Bla04 and PNGM-1_Q50_lddt*_ increased to 0.95 with seqID 26% over the aligned region. The active was captured as *high-lDDT* residues between these two structures, suggesting a conserved structural and functional core (Figure 5a).

Expression of *phnP* in *E. coli* has not been previously associated with β-lactam activity, and the metal-binding site is actually optimised for Mn^2+^ (as opposed to Zn^2+^) (Podzelinska et al., 2009). No signal peptide was detected in Bla04 using SignalP 6.0 (Teufel et al., 2022), suggesting the observed ampicillin activity potentially arose from mis-localisation, likely dependent on strong expression and/or indirect cytoplasmic buffering of the enzyme, rather than canonical periplasmic MBL activity. This is also supported by the lack of ampicillin observed in liquid media, where ampicillin diffusion is increased compared to solid media. Discrepancies between liquid and solid media have been reported with low agreement in *E. coli* for ampicillin (20%) (Palladini et al., 2023), as well as growth observed on agar, however not in LB (Schoning, 2014). The low copy number of the pGDP series may have also contributed to these effects (Cox et al., 2017; Wang & Joffré, 2025). Finally, no signal peptide was also detected using SignalP 6.0 in PNGM-1. It has been suggested the additional rRNase Z activity of PNGM-1 may point to the origin of class B3 MBLs (Lee et al., 2019), and *phnP* shares the tRNase Z endonuclease fold (Podzelinska et al., 2009).

## Discussion

Recent studies have demonstrated the success of *in silico* prediction of novel ARGs with high precision on testing datasets (Rannon et al., 2025; J. Wu et al., 2023), as well after experimental verification of ARGs with less than 30% homology to known ARGs (Ruppé et al., 2019). Nonetheless, computationally predicted novel ARGs that are clinically important (i.e., confer MICs at CLSI resistant breakpoints), more commonly have higher seqIDs to known ARGs (Alcock et al., 2023; Berglund et al., 2017, 2019; Lund et al., 2023; Marathe et al., 2019), and false positive predictions increase at lower seqIDs. Furthermore, high-throughput functional verification methods remain limited and continue to represent the principal bottleneck in the ARG discovery pipeline. To address this, we leveraged recent advances in computational protein structure prediction to distinguish functional ARGs from proto-resistance genes lacking key amino acid residues and localised structural features associated with antibiotic resistance. This approach prioritised conserved structural regions of known ARGs and the residues that code for them. By adopting automated MIC testing where possible, experimental validation confirmed the method as a novel and precise strategy for ARG discovery.

Nine out of nine potential ARGs identified from HMP reference strains by ARG-PASS were functionally verified to hold activity against streptomycin, trimethoprim, or β-lactams. Five proteins from HMP reference strains predicted non-functional were also experimentally confirmed to lack a resistance function, suggesting a low FNR for potential AAC(6’), *sul*, and class B1 β-lactamases. In addition, a *phnP* gene with only 17.3% seqID shared to known MBLs was functionally predicted from the AFDB. In total, eight out of 10 (80%) genes conferred an MIC at or above CLSI resistant breakpoints, with an average seqID of 46.5% to known ARGs.

The high precision of ARG-PASS may stem from its use of the superposition-free metric (lDDT) employed in FoldMason, to identify structurally conserved regions. Subsequent calculation of the pairwise TM-score vs seqID across these structurally conserved regions, means the algorithm captures the congruence of the overall (conserved) fold for use in downstream machine learning prediction. Inferring function by structural and sequence comparison of distantly modelled residues by restricting alignments to structurally conserved regions (*high-lDDT*), reduces noise from disordered and/or structurally divergent regions, which can hamper full protein structure alignments (Illergård et al., 2009; Koehler Leman et al., 2023). Clustering ARP structures at 20%, 30%, and 50% seqID before MSTA’s is thus a critical component of the method as it limits pairwise seqID and TM-score distances between *high-lDDT* ARP structures.

Subsequent assignment and comparison of a query protein’s TM-score and seqID features to a cluster–depending on its existing homology to known ARP structures–contributes to algorithmic precision at a range of seqIDs. However, ARP structures which don’t cluster due to distant homology (i.e., less than 20% homology) can limit direct comparison of qARP structure features to these ARPs. This limitation was addressed by relaxing ARG-PASS by omitting prior clustering, enabling the identification and functional confirmation of a highly divergent protein sharing only 17.3% sequence identity with known MBLs. We anticipate this limitation will be minimised as models are updated with ARGs identified through complementary approaches (for e.g., functional metagenomics, other computational tools).

The main required input to ARG-PASS is a qARP structure. Alignment of the qARP structure to *high-lDDT* ARP structures within each cluster and subsequent prediction by the one-class SVM, are designed for computational efficiency and high accessibility, and are completed in under a minute on a standard desktop or laptop computer. For qARPs of ∼300 amino acids in length, structure predictions can be completed in under 5 minutes on a GPU and under 20 hours on 8 CPUs. The AFDB can also be queried for a vast amount of protein sequences already with computational solutions, reducing the need for modelling a protein structure.

Although limited to six HMP reference strains, this study identified nine novel ARGs from the human gut, supporting the view that this microbiome serves as a reservoir of resistance (and pre-resistance) genes (Salyers et al., 2004). Although they appear to be chromosomally encoded, which aligns with findings from Ruppé et al. (2019), horizontal transfer to pathogenic strains remains possible, particularly under intense or prolonged antibiotic exposure. Two genes did not confer MICs at CLSI resistant breakpoints after expression in *E. coli*; however their stable integration within human microbiome-resident strains increases the risk of accumulating mutations and/or transferring to human pathogens, where clinically relevant MICs may be conferred. Finally, a novel streptomycin-resistant APH(6’) gene was identified from *P. aeruginosa* and *folA* genes from *P. aeruginosa, K. pneumoniae* subsp. *pneumoniae*, and *E. coli,* all provided TMP-resistance when expressed in *E. coli*. These species have known pathogenic strains, which increases their compatibility and spread in these hosts.

## Conclusions

The computational prediction and functional confirmation of novel ARGs in this study provides a proof-of-concept for training a one-class SVM on pairwise primary vs tertiary protein structure similarities in conserved regions of ARP structures. High-precision algorithms are essential for identifying novel ARGs while minimising false positives revealed by antibiotic susceptibility testing. In this study, this was achieved through rigorous experimental verification of both functional and non-functional predictions, using a standardised expression system and a range of susceptibility testing methods. Comparison of MICs to CLSI resistant breakpoints also delineated pre-resistance from resistance. Future large-scale investigations using ARG-PASS on human and environmental microbiome data may paint a more complete picture of numbers of potentially clinically relevant ARGs circulating these environments, particularly those with low homology to existing ARGs, as well as genes not previously associated with resistance. Finally, we suggest our approach can be expanded to other ARG classes as well as beyond predicting ARG functionality, and encourage researchers to experiment with other protein datasets.

## Methods

### Training set

#### Subject antibiotic resistance protein (ARP) structures

ARP structures of well-known ARG classes obtained from the CARD (Alcock et al., 2023) (AAC, APH, ANT, *dfr*, and *sul*, class A, B, C, and D β-lactamases, and PBP1 - PBP3’s) were downloaded from the AFDB (highest available model quality), or if unavailable predicted by AlphaFold2 (AF2) (Jumper et al., 2021), using full databases from 2023-05-30, or the AlphaFold3 (AF3) web server (Abramson et al., 2024). AF2 predictions were performed using Digital Research Alliance of Canada clusters or the COSMIC2 gateway web server (Cianfrocco et al., 2017). For PBP1-PBP3’s, which confer resistance by point mutations, the WT sequence was used as a representative sequence for protein structures because individual mutations are not expected to have major structural impact. Due to the high number of class A, C, and D β-lactamases (>400) which comprise highly similar variants, redundancy was reduced by clustering at 99% seqID over 99% length (of both query and subject) using Foldseek (v.10-941cd33) (van Kempen et al., 2024). This resulted in a total of 1172 ARP structures for use in creation of one-class SVM models. AlphaFold/UniProt accessions of ARP structures are presented in Additional file: Table S6.

#### Clustering of subject ARP structures

ARP structures were first sorted into classes (i.e., AAC, APH, ANT, dfr, and sul, class A, B1/B2, B3, C and D β-lactamases, and PBP’s) and clustered by Foldseek under default settings at 20%, 30%, and 50% seqID over any length. Clustering of ARP structures at seqIDs of 30% and 50% were deemed appropriate to group functional ARG homologues and are performed elsewhere to reduce functional redundancy (Mirdita et al., 2017; Suzek et al., 2007). To achieve a more inclusive threshold, 20% seqID was introduced to account for the high sequence divergence across ARG classes and to ensure most ARGs were clustered for downstream analysis. However, any clusters with less than 3 ARP structures were omitted from subsequent analysis.

#### Structurally conserved regions of ARP structures

To identify structurally conserved regions within clusters of ARP structures, we employed FoldMason (v.2-7bd21ed) (Gilchrist et al., 2026), a tool designed for structural alignment and consensus modelling of protein structures at scale. This release was current at the time of pipeline development; subsequent releases introduced changes to the alignment algorithm and lDDT scoring that reduced accuracy on the PCM dataset used for validation (Additional file: Table S5), and this release is therefore retained for reproducibility. FoldMason utilises a per-column local distance difference test (lDDT) score (Mariani et al., 2013) to assess the consistency of atomic positions across aligned structures. The lDDT can estimate local structural conservation across multiple structural alignments (MSTAs) without relying on superposition, and enables identification of conserved structural regions even in protein structures with global conformational variability. There are limited established lDDT score thresholds that correspond to conserved structural columns within MSTAs, and they are often context-dependent. In this study, we analysed the distribution of lDDT scores (excluding values of -1, where only one protein structure is aligned in the column) and extracted residues ≥ the upper 50th percentile (Q50), the upper 75th percentile (Q75), or the upper 90th percentile (Q90), depending on the number of protein structures in the MSTA (Table 2).

**Table 2.** lDDT threshold used to extract residues from ARP structures in MSTA to create high-lDDT ARP structures.

| Number of structures in MSTA | IDDT threshold | Example notation used for *high-IDDT* structure |
| --- | --- | --- |
| 0 - 25 | upper 50th percentile (median) | AAC(6')la<sub>Q50_iddt</sub> |
| > 25 - 200 | upper 75th percentile | dfrA17<sub>Q75_iddt</sub> |
| > 200 | upper 90th percentile | TEM-1<sub>Q90_iddt</sub> |

We found scaling the lDDT threshold with the number of structures in the MSTA was a useful trade-off between the identification of functionally important residues with the length of the *high-lDDT* region, and was a critical step for accurate pairwise seqID vs TM-score distributions.This also extracted structurally and functionally conserved residues within the context of the particular MSTA; as opposed to applying a pre-defined lDDT threshold across all MSTAs (which fluctuate in numbers of ARP structures and structural variability). We performed further validation of these metrics using an existing protein structure-based method called PCM, which published functional verification results of potential ARGs from various classes (see below).

After residue extraction, *high-lDDT* ARP structures were created, represented as new .pdb structure files for each ARP structure. The Foldseek *createdb* command was used to create databases of *high-lDDT* ARP structures within each cluster for downstream creation of one-class SVMs.

### Training

The training data represents pairwise (all-versus-all) primary and tertiary protein structure comparisons for *high-lDDT* ARP structures within 20%, 30%, and 50% seqID ARP structure clusters. In other words, an 𝑥, 𝑦 distribution of seqIDs (𝑥) vs TM-scores (𝑦) of *high-lDDT* ARP structures. This was computed using Foldseek TM-align on Foldseek databases created in the previous section with parameters --alignment-type 1 and --exhaustive-search specified, and TM-scores normalised by the subject length.

SeqID vs TM-score distributions were scaled using scikit-learn’s ‘StandardScaler’ (Pedregosa et al., 2011), which standardises each feature independently by removing its mean and scaling to unit variance. Parts of this workflow development were assisted using large language models (Anthropic and OpenAI), with outputs reviewed by authors.

#### One-class SVM and hyperparameters

One-class SVMs are semi-supervised machine learning algorithms commonly used for outlier detection (H. J. Shin et al., 2005) and were deemed suitable for this classification task due to their effectiveness in scenarios where negative labels are unavailable or unreliable. The one-class SVM algorithm learns a decision function that defines a boundary in a transformed feature space around the training data. This boundary is called the decision boundary (DB). New data points are classified as inliers or outliers based on their relative position to the DB (Schölkopf et al., 1999). For two-dimensional 𝑥, 𝑦 distributions, the shape and tightness of the DB are primarily controlled by the parameters ν (nu) and γ (gamma), and values for these parameters were automated to avoid manual adjustments. The parameter ∈(0,1] sets an upper bound on the fraction of outliers and a lower bound on the number of support vectors. Here, the training data represents a pairwise distribution of seqIDs (𝑥) vs TM-scores (𝑦) for *high-lDDT* ARP structures within clusters. One-class SVMs were trained on each distribution using scikit-learn (Pedregosa et al., 2011), and ν was selected as 1/𝑁, where 𝑁 is the number of training points and assumes that all training data are inliers. Ensuring the DB surrounds training points was important to reduce model bias from over-representation of variants with low variability (for e.g., TEM variants). The kernel parameter, γ, which defines the influence radius of a single data point in the radial basis function (RBF) kernel, was selected automatically using scikit-learn’s ‘scale’ option, which sets γ =1/(number of features × data variance), and provides a standard balance between overfitting and underfitting.

### Query antibiotic-resistant protein (qARP) structures for functional prediction

#### qARP structures from HMP reference strains

Protein coding sequences (CDS) were predicted from nucleotide sequences of whole genome sequences (WGS) originating from six Gram-negative HMP reference strains (The Human Microbiome Jumpstart Reference Strains Consortium et al., 2010) isolated from the human gut (Additional file 1: Table S1) by Prodigal v2.6.3 using default settings (Hyatt et al., 2010). We prioritised Gram-negative strains to increase the likelihood of successful expression in *E. coli* and included both fermentative and non-fermentative bacteria. CDS was then subjected to relaxed homology alignment using Diamond v2.1.16 (Buchfink et al., 2015) against a database we created of ARG sequences obtained from CARD v4.0.1 (Alcock et al., 2023). ARG sequences in this database match those used to create one-class SVM models. The following arguments specified were used for this Diamond screen: -e 10 --sensitive --query-cover 70 --subject-cover 70. Results for this screening against the HMP reference strains are presented in Additional file 1: Table S3. Protein structures of query ARGs not considered known ARGs (with seqID < 90%) were downloaded from the AFDB or predicted by AF2 or AF3 if unavailable, representing qARP structures for functional prediction by ARG-PASS. AlphaFold/UniProt accessions or computed AF2/AF3 structures of qARPs predicted functional and experimentally verified are listed in Additional file: Table S2.

#### Potential MBLs downloaded from the AFDB

To identify potential metallo-β-lactamases (MBLs) with low homology to existing MBLs and skip the initial Diamond screening against the custom CARD database, we obtained UniProt accessions from the UniProtKB database (The UniProt Consortium, 2025) with the following filter: (taxonomy_id:562) AND (protein_name:Metallo-beta-lactamase). qARP structures were then downloaded directly from the AFDB. qARP structures with >20% seqID to known MBLs were discarded and deduplicated by clustering at 99% seqID over 90% length (of both query and subject) using Foldseek, with the longest representative used as a qARP structure for functional prediction.

#### Cluster assignation

Before passing into one-class SVM models for functional prediction, qARP structures were aligned to all ARP structures of the ARG class, with Foldseek under default settings. This was performed so qARP structures could be assigned to 20%, 30%, or 50%, ARP structure clusters (cluster levels) for a particular ARG class, based on their highest seqID to an ARP structure(s) (Table 3). An approach which scaled with the homology of the qARP to known ARGs was deemed necessary to maintain algorithmic precision of functional predictions of qARPs at a range of seqIDs.

**Table 3.** Assignation of qARP structures to *high-lDDT* ARP structure cluster levels.

| Best seqID with ARP structure(s) (%) | Assigned to cluster level (seqID) |
| --- | --- |
| 0 - 25 | 20% |
| > 25 - 50 | 30% |
| > 50 | 50% |

As an exception, to maximise appropriate downstream functional predictions of the qARP to its closest ARP structure(s), if the qARP was assigned to a cluster level according to Table 3 but the ARP structure(s) for which it was assigned did not cluster at this level, it was dropped cluster levels until this was true. For example, qARP Aph01 had the highest seqID of 52.4% to the ARP structure encoded by APH(6)-Ic, so it was assigned to the 50% clusters. However, APH(6)-Ic clustered as a singleton at this level. It shares no other seqIDs >50% to other ARP structures, so Aph01 was dropped to the 30% cluster level. Again, the 4 ARP structures for which it shared >25-50% seqID did not cluster, so it was dropped to the 20% seqID cluster level and predicted by ARG-PASS.

### Cluster assignation omitted for potential MBLs from the AFDB

No qARP structures downloaded directly from the AFDB with less than 20% seqID to class B3 β-lactamases were predicted functional by ARG-PASS. Therefore, we excluded prior 20% seqID clustering specifically for class B3 MBLs, to create *high-lDDT* ARP structures across the entire B3 MBL class. qARP structures of potential MBLs downloaded directly from the AFDB were then assigned for functional prediction both to these *high-lDDT* class B3 MBLs, and to the 20% seqID cluster of *high-lDDT* class B1/B2 MBLs. All other steps of ARG-PASS remained the same.

We chose not to integrate the omission of prior clustering for highly divergent potential class B3 MBLs into the ARG-PASS pipeline to retain the high precision of the algorithm. Nonetheless, it demonstrates how reducing the clustering thresholds can relax ARG-PASS to identify more distant homologues, with the consideration that false positives may increase.

### Functional prediction

Each qARP structure was aligned using Foldseek to databases of clustered *high-lDDT* ARP structures, to obtain an 𝑥, 𝑦 distribution of qARP structure seqIDs (𝑥) vs TM-scores (𝑦) (also scaled using scikit-learn’s ‘StandardScaler’). As with the training data, this was computed using Foldseek’s TM-align mode, specified with --alignment-type 1, which uses the Needleman-Wunsch (NW) algorithm (Needleman & Wunsch, 1970) with affine gap costs. The NW algorithm enabled accurate global alignments of shorter *high-lDDT* ARP structures to full-length qARP structures, in contrast to Foldseek’s default alignment mode which uses the Smith-Waterman algorithm and is more suited to local alignments (Smith & Waterman, 1981). Additionally, Foldseeks’ --exhaustive-search was specified and the *high-lDDT* ARP structure (subject) length was used to normalise TM-scores and calculate seqIDs.

Let 𝑓(𝑥) be the decision function learned by the one-class SVM from the training data, where 𝑥 represents a 2D feature vector (i.e., seqID and TM-score). For a given qARP structure constituting multiple points 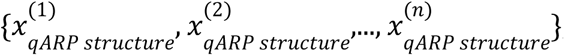, the qARP is classified as *functional* if *any* of its points lie on or within the DB:

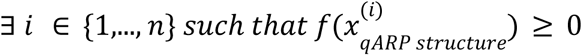

That is, the qARP is considered functional if at least one of its associated feature points has a decision score greater than zero, indicating it lies within the DB learned from the training data. Otherwise, the qARP is classified as non-functional. This criterion reflects the default decision threshold of the one-class SVM and assumes that any point outside the DB (i.e., 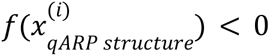) is an outlier, with respect to the training distribution.

#### Manual adjustment of Foldseek seqID calculations

Due to the alignment of shorter *high-lDDT* ARP structures against full-length qARP structures using Foldseek TM-align, seqID calculations were underestimated due to alignment gaps not being properly accounted for in the aligned region of the qARP structure. Manual adjustment of seqIDs was required by using an alignment length of the *high-lDDT* ARP structure (subject length).

### Validation of ARG-PASS on an existing protein structure-based method of predicting ARGs

Validation of ARG-PASS was performed on proteins previously experimentally verified by an existing computational method also based on protein structure used to identify ARGs, called pairwise comparative modelling (PCM) (Ruppé et al., 2019). PCM predicted ARGs from a 3.9 million gene catalogue of the intestinal microbiota. Referring to Supplementary Table 1 of the PCM paper, we collected protein sequences for analysis from the project website (https://mgps.eu/Mustard/index.php?id=accueil), and only focussed on ARG classes which have both positive and negative functional synthesis results (AAC(6’), APH, class A and B β-lactamases). Additionally, proteins with more than 80% seqID to known ARP structures were excluded. Protein structures were downloaded directly from the AFDB or predicted by AF2 and confirmed to be absent from the training distribution. Computational predictions and the functional synthesis results from the PCM paper, comparing ARG-PASS to PCM for proteins analysed are presented in Additional file 1: Table S5.

In summary, ARG-PASS correctly predicted 26 out of 33 proteins with an accuracy of 0.76. The precision increased from 0.64 across both ‘fair’ and ‘good’ predictions of PCM to 0.81 (Table 4). Furthermore, 13 out of 16 (0.81) true positive predictions were correct at seqIDs to a known ARP structure below 31%, compared to PCM (0.62) (Table 4).

**Table 4.**
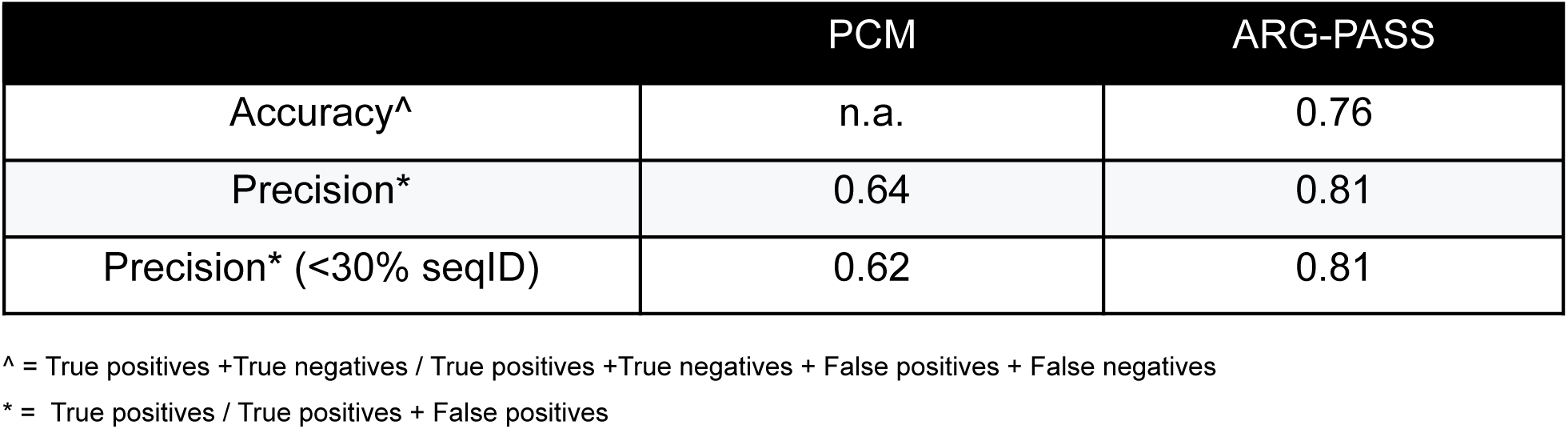
Accuracy and precision of PCM compared to ARG-PASS.

|  | PCM | ARG-PASS |
| --- | --- | --- |
| Accuracy<sup>^</sup> | n.a. | 0.76 |
| Precision<sup>*</sup> | 0.64 | 0.81 |
| Precision<sup>*</sup> (<30% seqID) | 0.62 | 0.81 |

<sup>^</sup> = True positives + True negatives / True positives + True negatives + False positives + False negatives
<sup>\*</sup> = True positives / True positives + False positives

It should be noted that ARG-PASS identified four false negatives (all at seqIDs below 25%). ARG-PASS was developed using ARG sequences strictly from the CARD, where resistance determinants are well-characterised and peer-reviewed (Alcock et al., 2023). As a result, most ARGs in the CARD likely confer clinically relevant MICs. This may have contributed to the identification of false negatives by ARG-PASS and verifying MICs of proteins from PCM within the experimental verification pipeline in this paper would offer a more coherent comparison. Furthermore, we found reducing the lDDT threshold for the extraction of structurally conserved residues (*high-lDDT* regions) of ARP structures generally decreased the FNR against the PCM dataset, however at the cost of algorithmic precision.

### Signatures of mobility near functional qARPs

To infer if functionally verified qARPs co-localised with MGEs, MGE-associated sequences (transposon, plasmid, phage, and integron-associated sequences) were queried 10 ORFs up and downstream of qARPs, by alignment against the MobileOG-db (Brown et al., 2022) using Diamond with a minimum seqID of 80% (over 90% query and subject length). In addition, the Galaxy server of PlasmidHunter (Tian et al., 2024) was used to infer if contigs containing qARPs had a chromosomal or plasmid origin.

### Experimental verification

The Minimal Antibiotic Resistance Platform includes vectors grouped by a kanamycin (pGDP1 and pGDP2) or ampicillin (pGDP3 and pGDP4) selective marker, and by promoter strength, with the stronger β-lactamase *bla* promoter (pGDP1 and pGDP3) and the weaker *lac* promoter (pGDP2 and pGDP4) (Cox et al., 2017). After codon optimisation for *E. coli*, DNA sequences of selected qARPs, positive controls not already available in the Minimal Antibiotic Resistance Platform, *dfrK* (NCBI accession CBL80435.1) and *sul3* (NCBI accession ACJ63260.1), and negative control *folA* from *E.coli* MG1655 (NCBI accession NP_414590.1) were synthesised by IDT. Note that no positive control was included for PBPs, as multiple attempts weren’t successful in assembling the β-lactam-resistant PBP3 from *E. coli* (CARD ARO: 3007423), into pGDP1 or pGDP2 of the Minimal Antibiotic Resistance Platform.

Synthesised gblocks were then assembled into appropriate pGDP vectors using the NEBuilder® HiFi DNA Assembly (New England Biolabs), transformed into *E. coli* MM294 by the chemical transformation method, and grown in Lysogeny Broth (LB) for 1h at 37°C with agitation. Colonies were grown overnight (o/n) on LB and agar plates containing 100 µg/mL of ampicillin or 50 µg/mL of kanamycin (selection). Transformants were recovered and grown o/n in LB with selection. Plasmids were then extracted using the Monarch Plasmid DNA miniprep kit (New England Biolabs) and sequenced on a minION Flongle Flowcell (Oxford Nanopore Technologies) to confirm an error-free assembly. Confirmed plasmids were then re-transformed via electroporation into *E. coli* BW25113 (Δ*bamB*Δ*tolC*), with the same subsequent steps as the chemical transformation method and stored at -80°C in glycerol. The Minimal Antibiotic Resistance Platform was a generous gift from Prof. Gerard Wright of McMaster University (also available as an Addgene kit #1000000143).

### MIC measurement

#### Broth microdilution (BM)

Determination of MICs by broth microdilution (BM) were measured in accordance with CLSI M07 (CLSI, 2024b). Cultures were grown overnight in MHII broth for 18h at 37°C, serially diluted, and applied in duplicate to microdilution plates containing MHII broth and appropriate antibiotics ranging around CLSI resistant breakpoints (CLSI, 2024a).

An approximate final concentration of 5 × 10⁴ colony forming units (CFU) was present in each well. Following incubation for 18h at 37°C, MICs were recorded as the antibiotic concentration that no visible growth was present, which was also confirmed by measuring the optical density at 600nm (OD600) of the well compared to wells without inoculation. All MICs were measured alongside previously transformed *E. coli* BW25113 (Δ*bamB*Δ*tolC*) strains containing an empty pGDP vector (negative control), and a pGDP vector containing a known ARG in the respective class of the qARP (positive control). A qARP was considered a functional ARG if its MIC was higher than that of the negative control in both duplicates.

### Spot plating assay (SPA)

The spot plating assay (SPA) was developed in-house in an attempt to increase through-put and automation of MIC measurements. Briefly, 2 µL of culture was collected by a Rotor HDA (Singer Instruments) from each well of a 96-well plate containing a strain for MIC testing, previously grown for 18h at 37°C in 200 µL LB, without selection. The 2 µL of culture was then spotted onto PlusPlates (Singer Instruments) containing MHII agar or MHII agarose (for SXT and TMP testing) and concentrations of appropriate antibiotics at CLSI breakpoints, in the same relative position as they were collected from the 96-well plate. After 18h at 37°C, plates were visually inspected, with an MIC recorded as the concentration for which no visible growth was present. A qARP was considered a functional ARG if its MIC was higher than that of the negative control.

### Phylogenetics

To investigate evolutionary relationships between functionally confirmed qARPs in this study and ARPs encoded by closely related ARGs, FoldMason under default settings was used to create an MSTA. Maximum likelihood was applied as a search criterion in IQ-TREE v.3.0.0 using the best-fit model automatically selected by ModelFinder (Nguyen et al., 2015). An Ultrafast Bootstrap analysis was performed with 1000 ML replicates and an approximate likelihood ratio test search with 10,000 replicates. Phylogenetic trees were visualised and edited with FigTree v.1.4.4 (available http://tree.bio.ed.ac.uk/software/figtree/).

## Supporting information

Additional file 2: Protein sequences of qARPs

Additional file 3: Structural alignments

## Declarations

### Ethics approval and consent to participate

Not applicable.

### Consent for publication

Not applicable.

### Availability of data and material

ARP and *high-lDDT* ARP structure databases, qARP structures analysed in this study, and reproducibility pipelines using ARG-PASS are available for download at 10.5281/zenodo.18817275. An example of the ARG-PASS method and code generated is also available at https://github.com/btroppo/ARG-PASS.

### Authors’ contributions

LB and LPH conceived and designed the study. LB and EBL performed PCR reactions, cloning assemblies, transformations, and functional verification of novel genes. FG performed Nanopore sequencing, provided experimental guidance and manuscript review. SR and LPH contributed to conceptual analysis, provided experimental guidance and manuscript review. LB designed the computational method, performed bioinformatic analyses, and drafted the manuscript. All authors read and approved the final manuscript.

## Acknowledgements

The authors would like to thank Professor Jean-Philippe Côté and Lya Blais (Department of Biology, Université de Sherbrooke, QC, Canada) for MIC testing using the Rotor HDA. The authors would also like to thank Professor Joseph Bielawski (Department of Biology, Dalhousie University, NS, Canada) for helpful discussions. This research was enabled in part by computational support provided by Calcul Québec (calculquebec.ca) and the Digital Research Alliance of Canada (alliancecan.ca).

