## Additional file 4: Figures S1 - S3 for "Conserved protein sequence-structure signatures identify antibiotic resistance genes from the human microbiome"

### Supplementary Figures

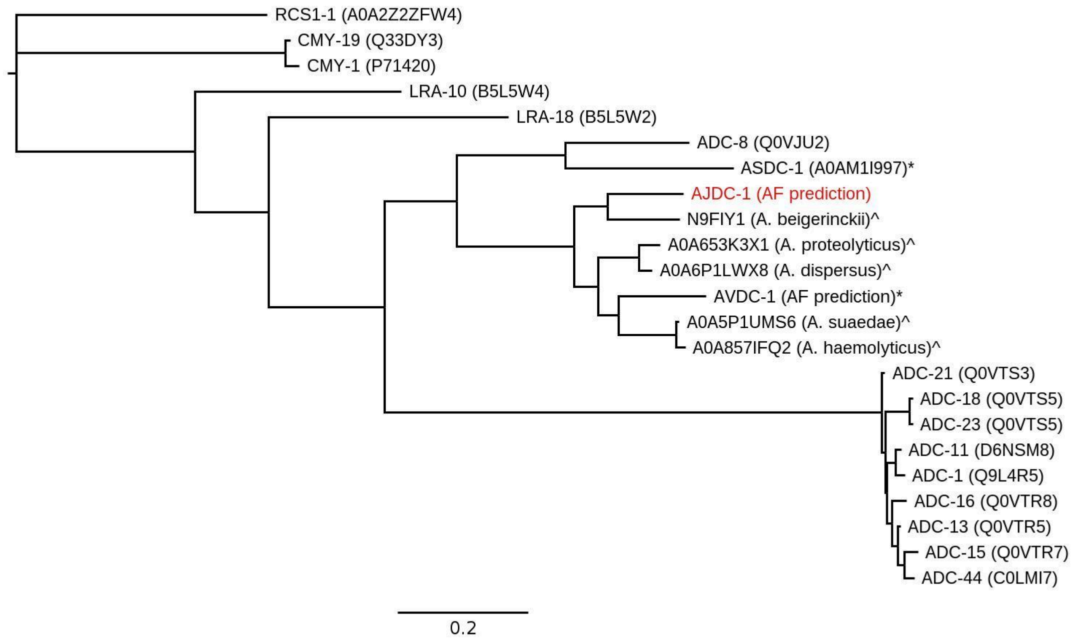


**Supplementary Figure 1.** Structural phylogeny of AJDC-1 (in red) with closest related known cephalosporinases. * = Cephalosporinases from *Acinetobacter* sp. in the BLDB and not in the CARD. ^ = Potential cephalosporinases from *Acinetobacter* sp. without experimental data, depicted with the host name.

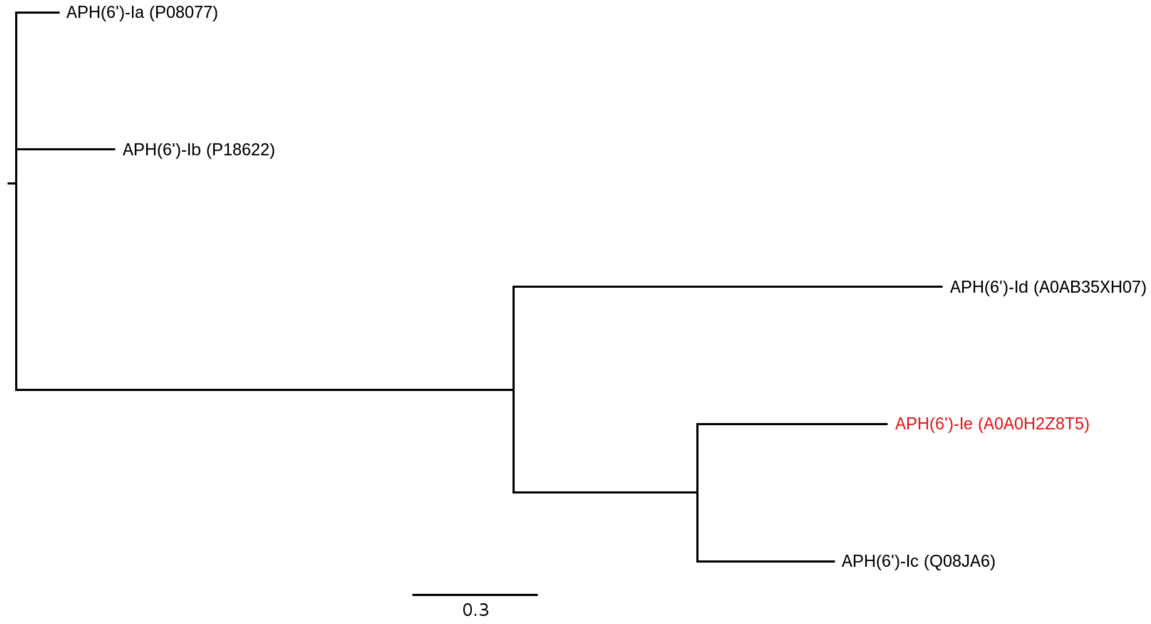


**Supplementary Figure 2.** Structural phylogeny of APH(6')Ie (in red) with remaining APH(6') genes from the CARD.

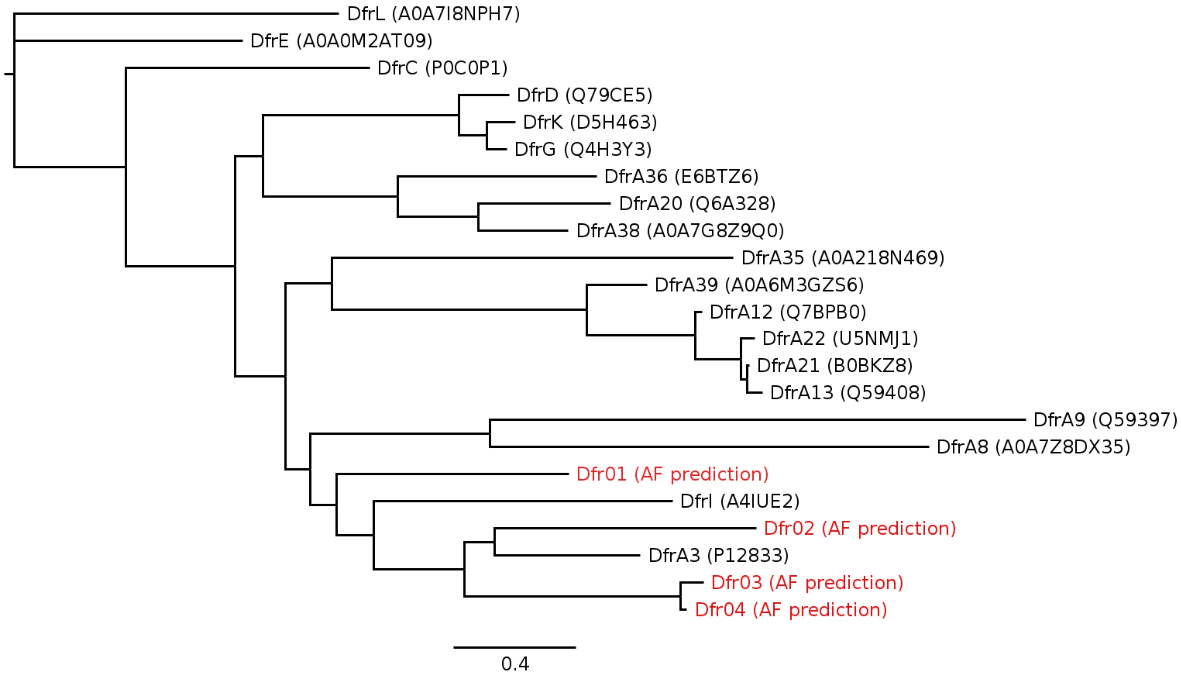


**Supplementary Figure 3.** Structural phylogeny of Dfr01-Dfr04 (in red) with remaining mobile Dfr variants within the one-class SVM cluster.
